# Single-cell profiling of the lung immune cells of diabetes-tuberculosis comorbidity reveals reduced type-II interferon and elevated Th17 responses

**DOI:** 10.1101/2024.10.24.619799

**Authors:** Shweta Chaudhary, Mothe Sravya, Falak Pahwa, V Sureshkumar, Prateek Singh, Shivam Chaturvedi, Debasisa Mohanty, Debasis Dash, Ranjan Kumar Nanda

**Affiliations:** Translational Health Group, International Centre for Genetic Engineering and Biotechnology, Aruna Asaf Ali Marg, New Delhi; National Institute of Immunology, Aruna Asaf Ali Marg, New Delhi; CSIR-Institute of Genomics and Integrative Biology, Mathura Road Campus, New Delhi; Bio-experimentational Facility, International Centre for Genetic Engineering and Biotechnology, Aruna Asaf Ali Marg, New Delhi; Institute of Life Sciences, Nalco Nagar Rd, NALCO Square, NALCO Nagar, Chandrasekharpur, Bhubaneswar, Odisha

**Keywords:** Diabetes, tuberculosis, scRNA sequencing, immunology, inflammation, T-cell immunity

## Abstract

Understanding the perturbed lung immune cells distribution and its functionality in tuberculosis (TB) is well documented; however, limited reports have covered their disruption, if any, in diabetes-tuberculosis (DM-TB) comorbid conditions. Here, we employed single-cell RNA-seq to investigate the molecular mechanisms that govern the heterogeneity in host immune response in DM-TB comorbid conditions. Diabetes is associated with chronic hyperinflammation and reduced lung-infiltrating immune cells, which delays the immune response to Mycobacterial infection. scRNA-seq of lung CD3^+^ and CD11c^+^ cells revealed compromised adaptive and innate immunity, with decreased Th1 and M1 macrophage populations in DM-TB mice. A dampened immune response, marked by increased IL-16 signaling and reduced TNF and IFN-II responses, was observed in DM-TB. This study highlights chronic inflammation, hyperglycemia, and dyslipidemia associated with diabetes impairing anti-TB immunity. Selective inhibition of aberrant IL-16 secretion and Th17 cell activation might provide strategies for better managing DM-TB comorbidity.

## Introduction

Tuberculosis (TB) is caused by the infection of *Mycobacterium tuberculosis* (Mtb) complex. The five major risk factors for TB include HIV infection, malnutrition, alcohol use disorders, smoking, and diabetes (Bagcchi, 2023; Pai et al., 2016). In 2021, globally one in every ten adults (537 million) were living with diabetes mellitus (DM) (Magliano and Boyko, 2021). DM patients show three times (95% CI, 2.27-4.46) higher risk of developing active TB disease, with six times higher odds of death during treatment (Dooley and Chaisson, 2009). Diabetic patients with TB (DM-TB) present a higher bacterial burden at case presentation and increased cavitary lesions (Dooley and Chaisson, 2009). Inferences from the euglycemic animal models and TB patients might have limited translatability in immunocompromised DM-TB patients. Therefore, it is crucial to understand the immunological interplay between DM and TB, which confers higher susceptibility, to develop and aid in effective management strategies for DM-TB comorbid conditions.

Higher baseline IL-6 in addition to increased type 1 and type 17 cytokines, was reported in DM patients (Gupte et al., 2022; Kumar Nathella and Babu, 2017). A delayed Mtb dissemination in the DM-TB mice model was reported with a slow rise of circulatory IFN-γ levels and a low CD14 and MARCO expression in alveolar macrophages (AM) (Cheekatla et al., 2016; Martinez and Kornfeld, 2014; Martinez et al., 2023; Vallerskog, Martens, and Kornfeld, 2010). The immune landscape of Mtb infected mice and macaque models demonstrated complex and extensive intracellular communication involving multiple cell types (Akter et al., 2022; Bromley et al., 2024; Gideon et al., 2022). Such detailed immune cell profiling studies in DM-TB condition, using Single-cell RNA sequencing (scRNA-seq) will bring better clarity then focusing on individual cell types like macrophages, NK or T-cells.

In this study, nicotinamide-streptozotocin (STZ-NA) based DM induction in C57BL/6 mice was attempted and aerosol-infected with low doses of Mtb H37Rv. At the peak of anti-TB adaptive immune response (Das et al., 2021), i.e., 21 days post-infection, in C57BL/6 mice, 10x Genomics-based scRNA-seq of lung CD3^+^ and CD11C^+^ cells along with serum cytokine profiling from DM-TB and control mice groups (DM, TB, and Healthy) were attempted to monitor immunological heterogeneity, if any. Higher basal circulatory IL-6, TNF-α, and IL-17A levels were observed in the DM groups irrespective of their Mtb infection status. scRNA-seq analysis of the lung tissue of DM showed sterile inflammation and inflammageing. The DM-TB group showed a delayed immune response, dampened pro-inflammatory Th1, M1 macrophage and CD4+CTL responses, and heightened Th17-mediated tissue pathology, which leads to higher bacterial load at the later time points. The observed lung immune cell heterogeneity between DM-TB and TB will be useful in designing therapeutics to better manage the DM-TB conditions.

## Results

### STZ-NA induced diabetes in mice mimics human T2DM complications

Patients with diabetes show a higher risk for respiratory infections such as influenza, COVID-19 and TB (Daryabor et al., 2020). However, the mechanism of this increased susceptibility at cellular and molecular levels needs focused studies. The current study aimed to understand the lung immunological landscape in diabetic and DM-TB comorbid conditions. We chemically induced type 2 diabetes mellitus (T2DM) following previous report with minor modifications (Cheekatla et al., 2016). Male C57BL/6 mice, 8–12 weeks old, received nicotinamide and streptozotocin (STZ-NA) intraperitoneal injection to develop DM and their blood glucose and change in body weight were monitored (Figure 1A). DM mice showed a significant decline in their body weight (Figure 1 B) and remained hyperglycemic throughout the experiment (Figure 1C). Intraperitoneal glucose tolerance test (ipGTT) and insulin tolerance test (ITT) of DM mice showed significantly higher insulin resistance (Figure 1D) and glucose intolerance (Figure 1E), compared to age and sex-matched controls. Dyslipidemia is another well-reported complication of T2DM and is a major risk factor for cardiovascular diseases. The DM mice showed significantly higher circulatory levels of triglycerides, total cholesterol, free fatty acids and significant hypoinsulinemia as compared to the euglycemic controls at 11 weeks post-induction (Figure 1F, G, H, I). Hyperglycemia, dyslipidemia, along with insulin resistance and glucose intolerance indicated that the STZ-NA model in C57BL/6 mice mimics T2DM and its associated complications.

**Figure 1:**
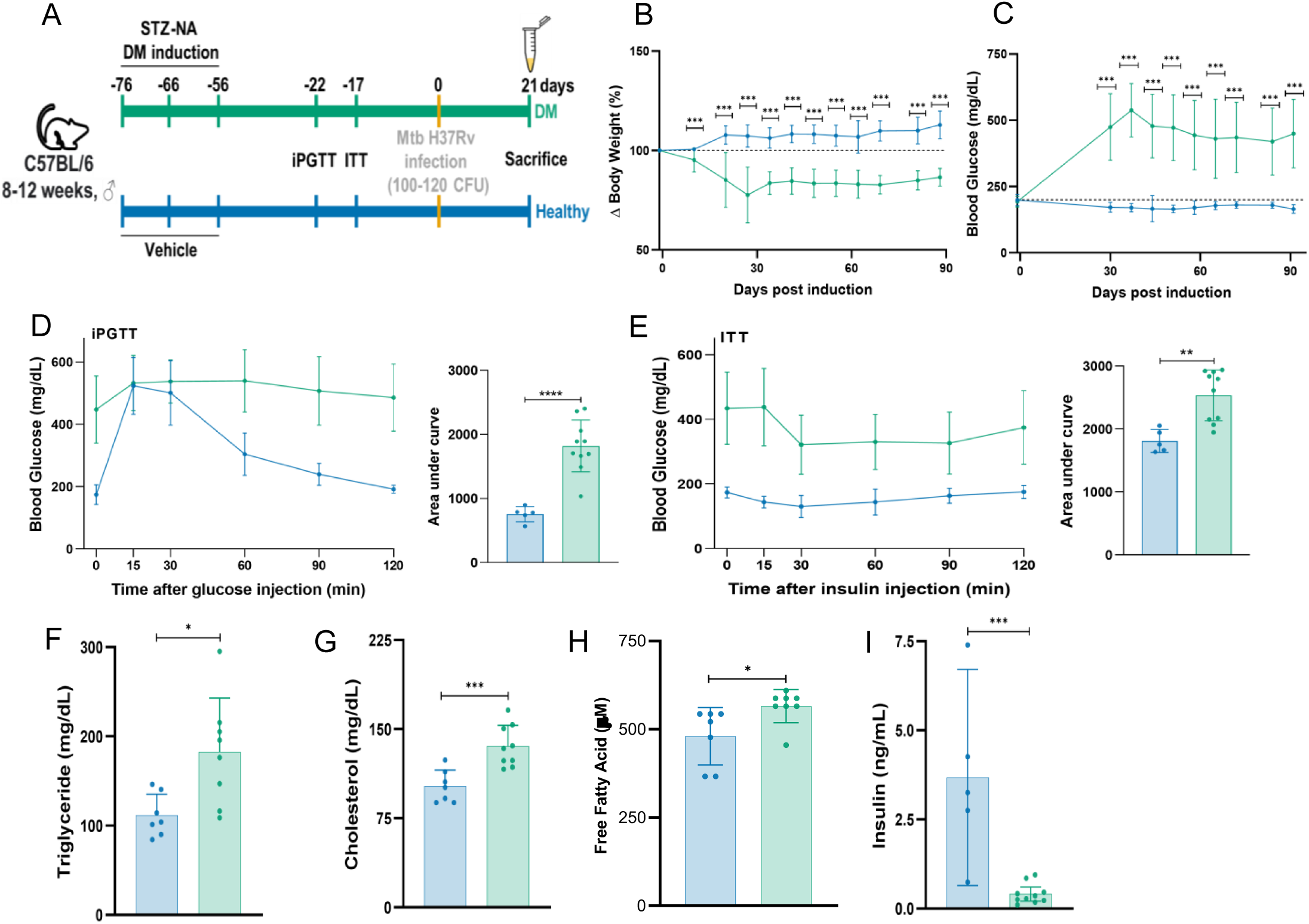
STZ-NA induced diabetes models Type 2 Diabetes Mellitus (DM) and its associated complications. (A) Schematic representation of the experimental plan. (B) Body weight change in DM and healthy mice during the course of the experiment. Healthy, n = 10; DM, n= 35. Error bars represent standard deviation. (C) Fasting blood glucose in DM mice and healthy controls during the experiment. Healthy, n = 10; DM, n= 35. Error bars represent standard deviation. (D) Insulin tolerance test (ITT) in DM and healthy controls. Healthy, n = 5; DM, n= 10. Error bars represent standard deviation. (E) Intraperitoneal glucose tolerance test (iPGTT) in DM and healthy controls. Healthy, n = 9; DM, n= 12. Error bars represent standard deviation. (F-H) Serum triglyceride, cholesterol, free fatty acid and insulin levels in DM mice and healthy controls. Healthy, n = 7; DM, n= 8. Error bars represent standard deviation. (I) Serum insulin levels in DM mice and healthy controls. Healthy, n = 5; DM, n= 10. Error bars represent standard deviation. Statistics: (B-I) unpaired t-test; ****: p < 0.0001, ***: p < 0.0005, **: p < 0.005, *: p < 0.05. A non-significant p-value was not indicated in the plots.

### Delayed immune response evident in DM-TB condition

These DM and healthy control mice were aerosol infected with a low dose (100-120 CFU) of Mtb H37Rv (Figure 2A). At 21 days post-infection (d.p.i), similar gross lung and spleen pathology between TB or DM-TB groups was observed (Figure 2B, 2C, 2E). However, the lung inflammation score, based on hematoxylin and eosin (H&E) staining, was highest for the DM-TB group (Figure 2D). Cellular infiltration to the lungs was evident in both groups at this time point. However, the inflammation was more dispersed in the lung tissue of the DM-TB group than in euglycemic control (Figure 2D, Supplementary Figure S1). At 21 d.p.i, lung and spleen bacterial burden was similar (Figure 2F) in the TB and DM-TB groups, corroborating earlier report (Cheekatla et al., 2016). However, serum pro-inflammatory cytokine levels, measured using LegendPlex^TM^, showed group-specific differences (Figure 2G, Supplementary Figure S2). IFN-γ is a type II interferon released by the Th and NK cells, activates macrophages to produce reactive nitrogen species to counteract microbial infection. IFN-γ also stimulates macrophages to overcome phagolysosome maturation by activating apoptosis via caspases 3 or 7, thus blocking mycobacterial replication (Shanmuganathan et al., 2022). By limiting excessive neutrophil migration to the site of infection, it prevents hyperinflammation in the lung (Shanmuganathan et al., 2022). Significantly higher circulatory IFN-γ levels were observed in both TB and DM-TB groups compared to the uninfected control, corroborating earlier reports (Domingo-Gonzalez et al., 2016) (Figure 2G). TNF-α initiates cytokine response against Mtb by inducing the CXCL9 and CXCL10 expression, which facilitates T-cell recruitment at the infection site (Robert and Miossec, 2021; Roca et al., 2022). TNF-α is associated with insulin resistance development in T2DM patients (Stephens, Lee, and Pilch, 1997). Higher circulatory pro-inflammatory cytokines such as TNF-α, IL-6 and IL-1β levels are reported in the T2DM patients and mice models (Boni et al., 2022; Cheekatla et al., 2016; Gupte et al., 2022). We observed significantly higher circulatory TNF-α levels in DM, TB and DM-TB conditions compared to healthy controls, while IL-6 and IL-17A levels were elevated in DM and DM-TB conditions (Figure 2G). Higher IL-6 levels in DM-TB damage the lung tissue (Cheekatla et al., 2016). Higher IL-6 levels in DM-TB comorbid patients before treatment are associated with worsened treatment outcomes (Gupte et al., 2022). Increased IL-17A levels in newly diagnosed T2DM patients have been reported, and the circulatory IL-17A levels positively correlate with TNF-α levels (Abdel-Moneim, Bakery, and Allam, 2018). In this study, significantly higher circulatory IL-6, IL-17A and TNF-α levels were observed in the TB and DM-TB groups, corroborating earlier reports (Domingo-Gonzalez et al., 2016).

**Figure 2:**
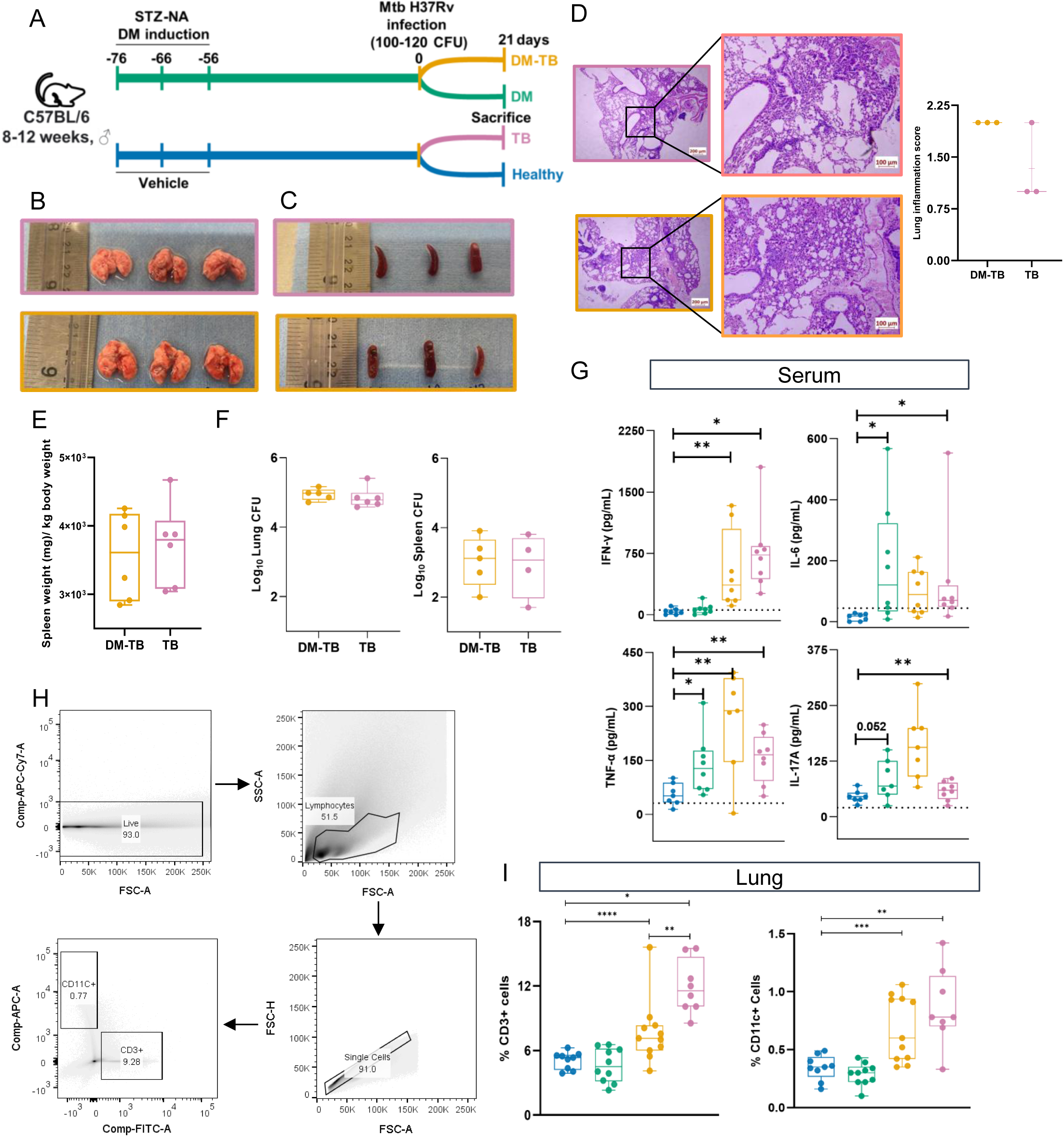
At 21 days post-Mtb H37Rv infection, hyperglycemic (DM) mice and DM-TB mice display comparable tissue bacterial loads to Mtb infected (TB) group with distinct serum cytokine and lung immune cell profiles. (A) Schematic representation of the experimental plan. (B, C) Gross lung and spleen pathology in Mtb H37Rv infected mice at 21 days post-infection (d.p.i.). (D) Hematoxylin and Eosin-stained lung tissue sections show histopathology differences and lung inflammation scores between groups. TB, n = 3; DM-TB, n = 3. Error bars represent standard deviation. (E) Spleen weight at 21 d.p.i. in DM-TB and TB mice. TB, n = 6; DM-TB, n = 6. Error bars represent standard deviation. (F) Lung and spleen bacterial burden at 21 d.p.i. in DM-TB and TB mice. TB, n = 6; DM-TB, n= 5 for lung CFU. TB, n = 4; DM-TB, n = 5 for spleen CFU. Error bars represent standard deviation. (G) Serum IFN-γ, Il-6, TNF-α and Il-17A levels (pg/mL) measured via LEGENDplex assay in healthy, DM, TB and DM-TB groups at 21 d.p.i. Healthy, n = 7; DM, n = 8; TB, n = 8; DM-TB, n = 8 for IFN-γ and IL-6. Healthy, n = 7; DM, n = 8; TB, n = 8; DM-TB, n = 7 for TNF-α and Il-17A. Error bars represent standard deviation. (H) The gating strategy was used to sort lung CD3+ and CD11C+ cells from all study groups. A representative sample from the TB group is used. (I) Frequency (%) CD3+ and (B) CD11C+ cells in healthy, DM, TB and DM-TB mice at 21 d.p.i. Healthy, n = 9; DM, n = 10; TB, n = 8; DM-TB, n = 11. Error bars represent standard deviation. Statistics: (D-I) unpaired t-test; ****: p < 0.0001, ***: p < 0.0005, **: p < 0.005, *: p < 0.05. A non-significant p-value was not indicated in the plots.

Alveolar macrophages present Mtb with a preferred niche, and the pathogen is known to delay antigen presentation by both macrophages and dendritic cells at the nearby draining lymph node. The immune response against Mtb starts in the draining lymph nodes and not in the lungs (Chackerian et al., 2002). Therefore, macrophages and dendritic cells are crucial in disseminating the pathogen to initiate host defence. T cells are well known for their anti-TB responses. DM mice show a three-day delay in Mtb dissemination than euglycemic controls, contributing to a compromised immune response (Vallerskog, Martens, and Kornfeld, 2010). As 21 days post Mtb infection is the peak of immune response in C57BL/6 mice, we focused on this crucial time point to study anti-TB response (Das et al., 2021).

From the lungs of TB, DM-TB, DM and healthy mice, CD11c^+^ (DCs, macrophages and resident monocytes), and CD3^+^ (T and NK cells) were flow-sorted (Figure 2H). Mtb H37Rv infection led to a significant increase in the lung CD11c^+^ and CD3^+^ populations compared to healthy control, the overall CD3^+^ cells frequency was significantly lower in the DM-TB compared the TB group (Figure 2I). A reduction in CD11c^+^ cells was observed in DM-TB mice compared to TB but was statistically insignificant (Figure 2I). At 21 d.p.i., the lung Mtb load was similar between DM-TB and TB groups, but a significant difference was observed at the immunological scale.

### Single-cell RNAseq profile of diabetic lung

For in-depth characterization of the lung immune cell landscape of DM mice and euglycemic healthy controls, paraformaldehyde-fixed samples were taken for scRNAseq experiments (Figure 3A). Following single-cell alignment and quality control steps, 5,364 and 7,838 cells from healthy and DM groups respectively, qualified the criteria (Supplementary Figure S3). Based on the canonical gene expression, 3 major cell types: T-cells (*Cd3d+*), B-cells (*Ms4a1+*) and myeloid cells (*Cd14+ Itgam+, Itgax+*) were identified (Figure 3B, Supplementary Table S1). Since B-cells were not selectively flow-sorted, the number was quite low, and excluded from further analysis. Between the healthy and DM groups, 14 cell types, with eight subtypes in the T cell cluster and six subtypes in the myeloid cell cluster, were identified. T cell subclusters included Naïve CD4+ T cells (Cd4+ Sell+ Ccr7+), Naïve CD8+ cells (Cd8a+ Sell+ Ccr7+), Th1 cells (Cd4+ Tbx21+ Ifng+), Th17 (Cd4+ Rorc+ Il17ra+ Il23r+), Treg (Cd4+ Foxp3+ Ctla4+), Tfh (Bcl6+ Maf+ Cxcr5+), CD4+ CTLs (Cd4+ Mki67+ Gzmk+ Gzma+) and CD160+ NKT cells (Cd3d+ Cd8a+ Nkg7+ Klrd1+) based on the expression of known markers. The lung myeloid cell compartment was composed of AMφ (Marco+ Siglecf+), pDC (Bst2+ Siglech+), mDC (Ifi30+ Flot1+ Psmb9+), cDC1 (Xcr1+ Cd24a+ Clec9a+), classical monocytes (Cd14+ Fcgr3-Sell+ Ccr2+) and intermediate monocytes (Cd14+ Fcgr3+ Treml4+ Sell+) based on the expression of known markers (Figure 3C, 3D, Supplementary Table S2). Interestingly, the overall lung Th17 cell proportion was higher in the DM mice than in healthy controls (Figure 3E, Supplementary Table S3). Higher Th17 cell numbers are implicated in inflammageing-associated chronic diseases such as T2DM (Abdel-Moneim, Bakery, and Allam, 2018). In addition, the number of naïve CD4+ T cells was higher in DM mice lungs than in healthy controls (Figure 3E). Thus the scRNAseq analysis of lung immune cells of diabetic mice revealed a dysregulated immune cell landscape characterized by an elevated frequency of inflammatory Th17 cells and naïve CD4+ T-cells.

**Figure 3:**
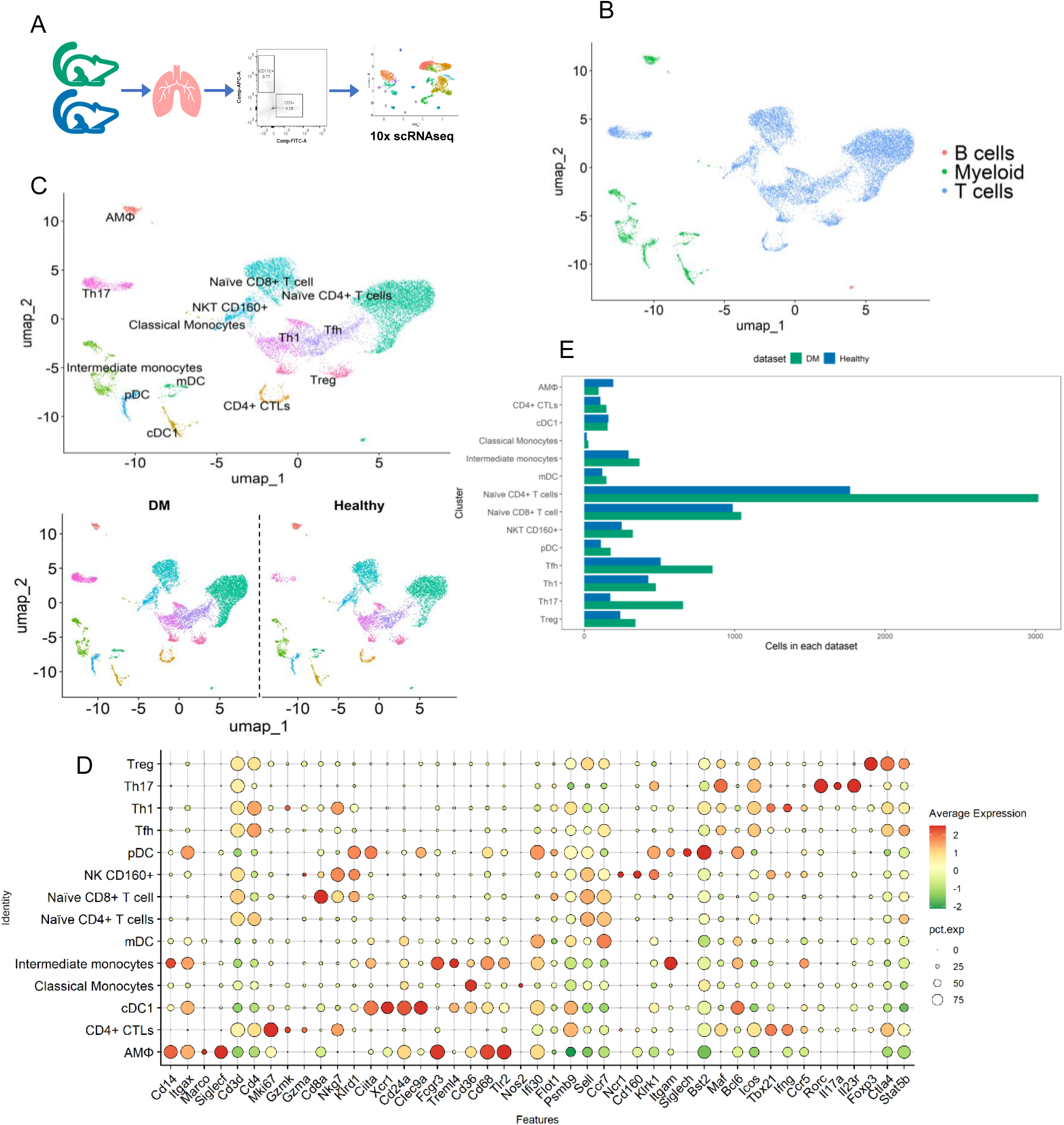
scRNA-seq analysis shows marked alterations in the lung immune cell profiles of diabetic mice. (A) Schematic representation of the experimental plan. (B) Uniform Manifold Approximation and Projection (UMAP) visualization of the major cell types representing lymphoid/myeloid cells in dataset. (C) Annotated UMAP showing lung immune cell subtypes in DM and Healthy mice. (D) Dot plot indicating expression of key genes detected for each cell subtype. (E) A stacked bar plot shows the percentage of cell subtypes in each group.

### T-cells of diabetic mice showed altered gene expression patterns

In the DM group, higher overall Tfh, Th1, Th17, Tregs, and naïve CD4+ T cells were observed compared to the healthy control (Figure 3E). DeSeq2 analysis of naïve CD4+ T-cells in DM group showed a set of 320 differentially expressed genes (DEGs, log_2_FC>±1.0, p<0.05; 88/232: up/down) (Figure 4A, Supplementary Table S4). Gene set enrichment analysis (GSEA) of the deregulated genes showed enrichment of type 1 interferon responses, inflammasome-mediated signalling, pyroptosis, IL-1 production, T-cell proliferation, and innate immune responses. NLRP3 inflammasome activation is associated with IL-1β release and pyroptosis (Supplementary Figure S4, Supplementary Table S5). Pyroptosis has been implicated in sterile inflammatory diseases, particularly in T2DM and lung inflammation in asthma (Li et al., 2021). Reports show that NLRP3 inflammasome-mediated pyroptosis contributed to islet inflammation in T2DM patients and chemically induced diabetic rat models (Li et al., 2021). This study shows that NLRP3-mediated pyroptosis is not limited to pancreatic islet cells and induces a sterile inflammatory state in the lungs of T2DM mice models. Type 1 interferon responses have an antagonistic effect on IL-1 responses, and IFN-β has been known to suppress IL-1 transcription and translation (Mayer-Barber and Yan, 2017). In the DM mice, IFN-β may be anti-inflammatory by limiting IL-1 driven immunopathology. The naïve T-cells in the DM group showed a depressed regulation of biochemical pathways, such as glucose, fatty acid, organic acid, prostaglandin transport, triglyceride, fructose-6-phosphate, glutamine, and long-chain fatty acyl CoA metabolism (Supplementary Figure S4). As these metabolic pathways support the energy demand and the overall metabolic flexibility of naïve T cells seems to be compromised in diabetic mice and show an enhanced inflammatory state.

**Figure 4:**
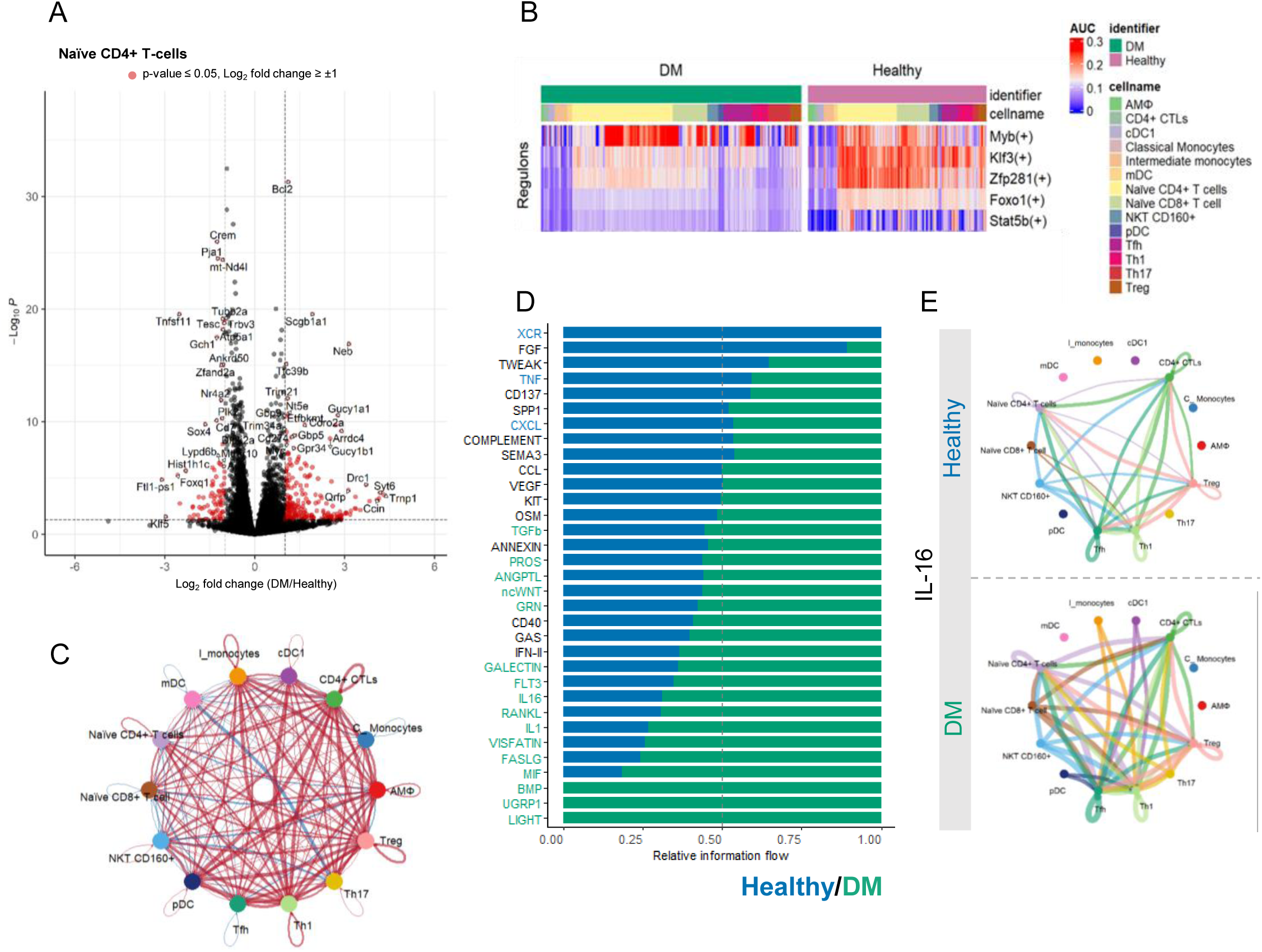
Altered functional, transcriptional, and intercellular communication profiles in lung immune cells of diabetic mice. (A) Volcano plot showing significantly up-/down-regulated genes in naïve CD4^+^ T-cells between DM and healthy mice. Significantly deregulated genes: p-value ≤ 0.05, log_2_fold change ≥ ± 1. (B) Heat map showing a subset of differentially activated regulon for cell clusters in hyperglycemic (DM) and euglycemic (healthy) groups. An unpaired Wilcoxon rank-sum test was performed to compare the activity of each regulon between datasets (healthy and DM). p-value ≤ 0.05 was selected for finding the significant difference in regulon activity. (C) CellChat circle plot showing signaling networks in DM versus healthy groups. The thickness of the edge represents the strength of signaling, and the red and blue colour represent upregulated and downregulated interaction, respectively, in the DM group compared to the healthy control. (D) A stacked bar plot showing the relative information flow between the ligand-receptor pairs in healthy and DM groups. A paired Wilcoxon test was performed to determine whether there was a significant difference in the signaling information flow between healthy and DM groups. The top signaling pathways coloured green are enriched in DM, and those coloured blue are enriched in the healthy group. p-value ≤ 0.05 was selected for finding the significant difference in regulon activity. (E) CellChat circle plot showing IL-16 signaling network in DM and healthy groups. The thickness of the edge represents the strength of signaling.

### Decoding lung immune cell transcription regulation in diabetes

Employing single-cell regulatory network inference and clustering (SCENIC) analysis dynamics of the transcription factors across cell types present in the lungs of DM mice were monitored (Figure 4B, Supplementary Figure S4, Supplementary Table S6). Interestingly, the Klf3, Zfp281 and Foxo1 transcription module activity was suppressed in the immune cells of the DM group (Figure 4B). Klf3 predominantly functions as a transcription repressor and is known to maintain T-cell quiescence. Klf3 knockout leads to myeloproliferative disorder and systemic inflammation in mice models (Cao et al., 2010). Lymphocytes namely, naïve CD4+ T-cells, naïve CD8+ T cells, Tfh, Th1, Th17, Tregs and NKT Cd160+ cells, showed a reduced Klf3 regulation in the DM group then the healthy controls, indicative of the activated state of these cells. Zfp281 regulates CD4+ T cell activation and cytokine production by repressing Ctla-4 transcription (Guo et al., 2020). While Zfp281 knockdown does not affect T-cell development, it has been reported to impair CD4+ T-cell activation and cytokine production in response to *Listeria monocytogenes* (Guo et al., 2020). A repressed Zfp281 regulon activity observed in the DM group compared to healthy controls (Figure 4B). Forkhead box O (Foxo) family of proteins are responsible for T-cell survival, trafficking, differentiation and memory response and their activity is inversely related to senescence (Delpoux et al., 2021). A diminished Foxo1 activity was observed in all lung T-cell subtypes (naïve CD4+, Naïve CD8+, Th1, Th17, Treg and Tfh cells) of the DM group, indicating impaired functionality (Figure 4B). Consequently, the reduced Klf3 expression in T-cells from the DM group indicates a heightened activation state and loss of quiescence, even without a pathogenic challenge. Meanwhile, reduced Foxo1 activity supports the inflammaging phenotype observed in the lungs of diabetic mice.

### Impact of hyperglycemia on pulmonary inflammation

In the DM group, the total number and strength of cell-cell communications was significantly high as inferred from the CellChat analysis compared to the healthy control (Figure 4C, Supplementary Figure S5). The cell signaling mediated by CD4+ CTLs, Tregs, Th1, naïve T-cell and intermediate monocytes was relatively increased in the DM lungs, and signals sent by mDCs, naïve CD8 T-cells and Tfh cells were decreased (Figure 4C). The deregulated ligand-receptor interactions were compared between DM and healthy control. MIF, IL16, FLT3 and LIGHT signaling pathways were significantly enriched in the lungs of DM mice (Figure 4D, Supplementary Table S7). The sender-receiver dynamics in healthy mice showed that the CD4+ CTLs were majorly responsible for MIF release, but in the DM group, both lymphoid (CD4+ CTLs, Naïve CD8+, NKT CD160+ and Th17 cells) and myeloid (alveolar macrophages) population were involved in sending MIF signals (Supplementary Figure S5). MIF is a lymphokine involved in immunoregulation, and the inflammation is mediated by its ability to promote pro-inflammatory cytokines such as TNF-α, IL-1β and IL-6. In addition, MIF also overrides the anti-inflammatory effects of glucocorticoids (Calandra and Roger, 2003; Joost J. Oppenheim, 2000).

IL-16 is a pro-inflammatory cytokine that works as a chemoattractant for CD4+ cells (T-cells, eosinophils and dendritic cells), prevents antigen-stimulated cell death and prime T-cell activity via IL-2. IL-16 has also been implicated in airway inflammation in the C57BL/6 model for asthma (Li et al., 2019a). We observed an increased IL-16-mediated cellular crosstalk in the DM group (Figure 4E). CD8+ T cells produce IL-16 in response to an antigenic stimulus, while CD4+ T cells have been reported to have a constitutive expression of pro-IL-16 molecule, which gets cleaved upon activation. We observed that Th17 and naïve CD8+ T-cell-based IL-16 interactions were unique to the DM condition. Similarly, cDC1, pDC, and intermediate monocyte-based interactions were also unique for the lungs of diabetic mice. The lymphocyte chemotactic ability of DCs has been reported to be IL-16 mediated. LIGHT is a member of the TNF superfamily and stimulates the growth and activation of T-cells. LIGHT has also been implicated in the pathology of T2DM and is responsible for attenuating insulin release, mediating vascular inflammation and regulating glucose metabolism via GLUT4 expression and signaling for this ligand was specifically enriched in the DM group only (Figure 4D, Supplementary Table S7) (Halvorsen et al., 2016; Kou et al., 2021). The enhanced signaling of MIF and IL-16 in the lungs of diabetic mice indicates a more pronounced basal inflammatory state within the system. The upregulation in pro-inflammatory signaling points to a chronic, underlying inflammation that may contribute to or exacerbate the overall immune dysfunction observed in diabetes.

### Immune cell dynamics in the lungs of diabetes-tuberculosis comorbid condition

A similar bioinformatics pipeline was employed to filter and integrate the TB and DM-TB data sets, resulting in 5,747 and 5,888 cells, respectively (Figure 5A, Supplementary Figure S3). Using previously discussed cell markers, three major cell types namely, T-cells, B-cells and myeloid cells were identified in the integrated TB and DM-TB dataset (Figure 5B, Supplementary Table S1). Upon further classification, we identified 15 cell types, with 8 subtypes in the T cell cluster (Naïve CD4+ T cells, Naïve CD8+ cells, Th1 cells, Th17, Treg, Tfh, CD4+ CTLs and CD8+ effector T cells) and 7 in the myeloid cell cluster (AMφ, pDC, mDC, cDC1, intermediate monocytes, NK Cd160+ and M1 macrophages) (Figure 5C, Supplementary Table S2, Figure 5D, Supplementary Figure S6, S7). Compared to TB only mice, overall frequencies of naïve CD4+, naïve CD8+, Th17 cells, and AMφ were higher in DM_TB mice, while frequencies of CD4+ CTLs, Th1 cells, intermediate monocytes, and M1 macrophages were lower (Figure 5E, Supplementary Table S8). Overall, scRNAseq analysis of the lung immune cells from *Mtb* infected diabetic (DM-TB) mice showed a dysregulated anti-TB response, characterized by increased naïve T-cells and reduced pro-inflammatory CD4+ CTLs, M1 M , and Th1 cells.

**Figure 5:**
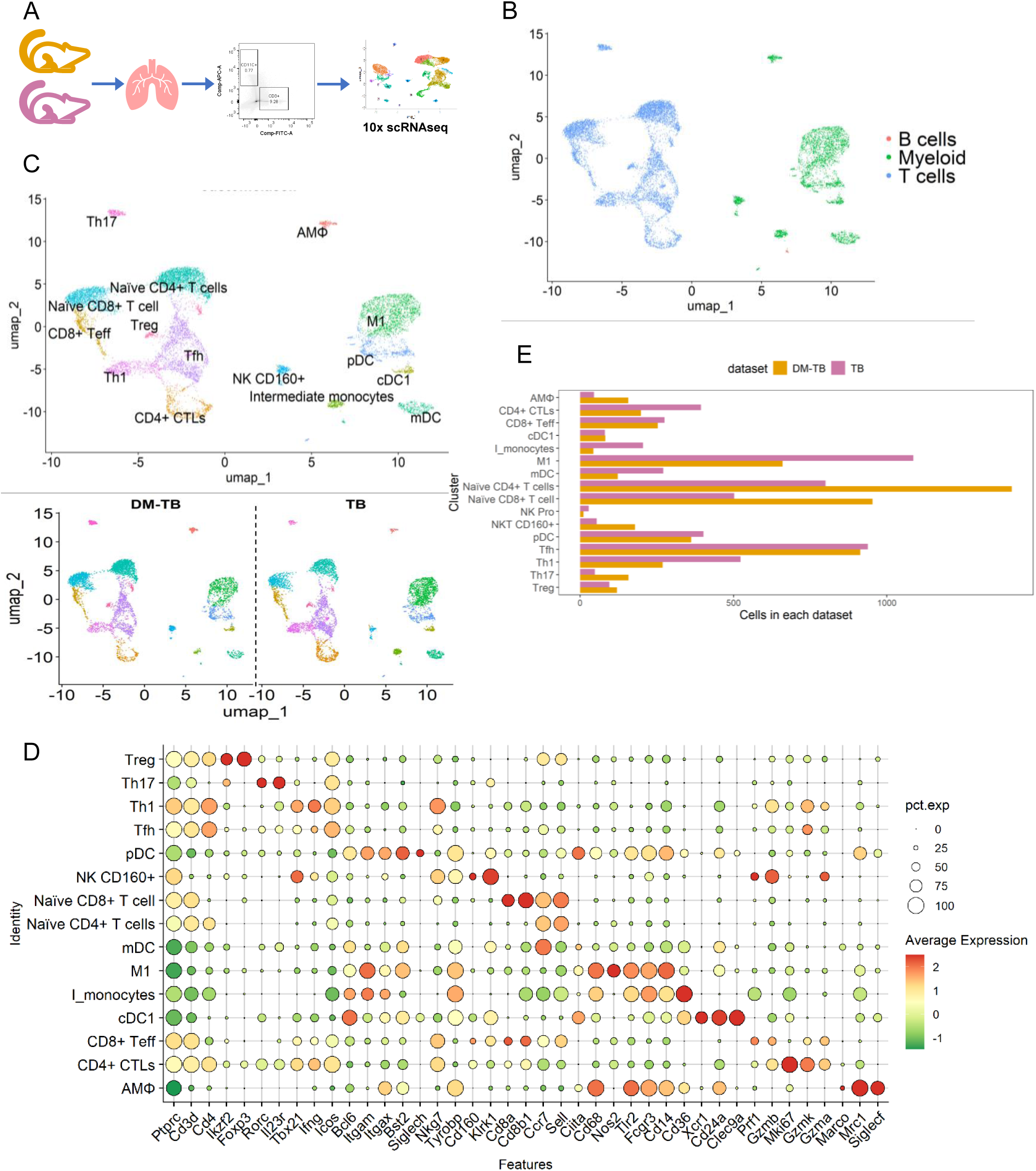
Impact of Mtb H37Rv infection on lung immune cell landscape of diabetic mice. (A) Schematic representation of the experimental plan. (B) Uniform Manifold Approximation and Projection (UMAP) visualization of the major cell types representing lymphoid/myeloid cells in the dataset. (C) Annotated UMAP showing lung immune cell subtypes in DM-TB and TB mice. (D) Dot plot indicating expression of key genes detected for each cell subtype. (E) A stacked bar plot shows the percentage of cell subtypes in each group.

### Differential gene expression highlights dysfunctional T-cells in the lungs of diabetes-tuberculosis comorbidity

In the DM-TB lungs, a low number of M1 macrophages, Th1 and CD4+ CTLs, while higher Th17, alveolar macrophages, naïve CD4+ and naïve CD8+ T cells were observed compared to the TB group. A set of 1,631 DEGs (922/709: down/up) in M1 macrophages of the DM-TB group was identified (Figure 6A, Supplementary Table S9, Supplementary Figure S10). GSEA analysis showed enrichment of nitric oxide (NO) mediated signal transduction, regulation of autophagy, response to glucocorticoid stimulus, negative regulation of cell-substrate adhesion, TGF-β signaling pathway and hematopoietic stem cell homeostasis (Supplementary Figure S10, Supplementary Table S10). The genes that regulate peptide cross-linking pathways included Thbs1, Stfa3, Fn1 and F13a1. Thbs1, Fn1 and F13a1 are involved in the modulation of extracellular matrix and are correlated with reduced macrophage inflammatory potential, further corroborated by the positive enrichment of TGFβ signalling.(Li et al., 2019b; Porrello et al., 2018; Stein et al., 2016; Xiao et al., 2018) M1 macrophages of the DM-TB group had significantly higher Trem2, Dapk1 and Bmf gene expression. While Trem2 and Dapk1 positively regulate autophagy, Bmf promotes cell survival, thereby negatively regulating autophagy. Negatively enriched GO terms for DM-TB M1 macrophages included responses to type-I and type-II interferons, AIM2 inflammasome complex assembly, TLR9 signaling pathway, regulation of Th1 responses, pathogen recognition receptor signaling, and innate (NK cell-mediated) as well as adaptive (B cell and Th1 cell-mediated) immune responses (Supplementary Figure S10). We also performed DEG analysis of Th17 and Tfh cells in the DM group compared with healthy controls. In Th17 cells from the DM group, we identified 294 DEGs, of which 109 genes were significantly downregulated and 186 genes were significantly upregulated compared with healthy controls, whereas 231 DEGs were identified in Tfh cells from DM lungs, with 67 genes significantly downregulated and 164 genes significantly upregulated (Supplementary figure S10).

**Figure 6:**
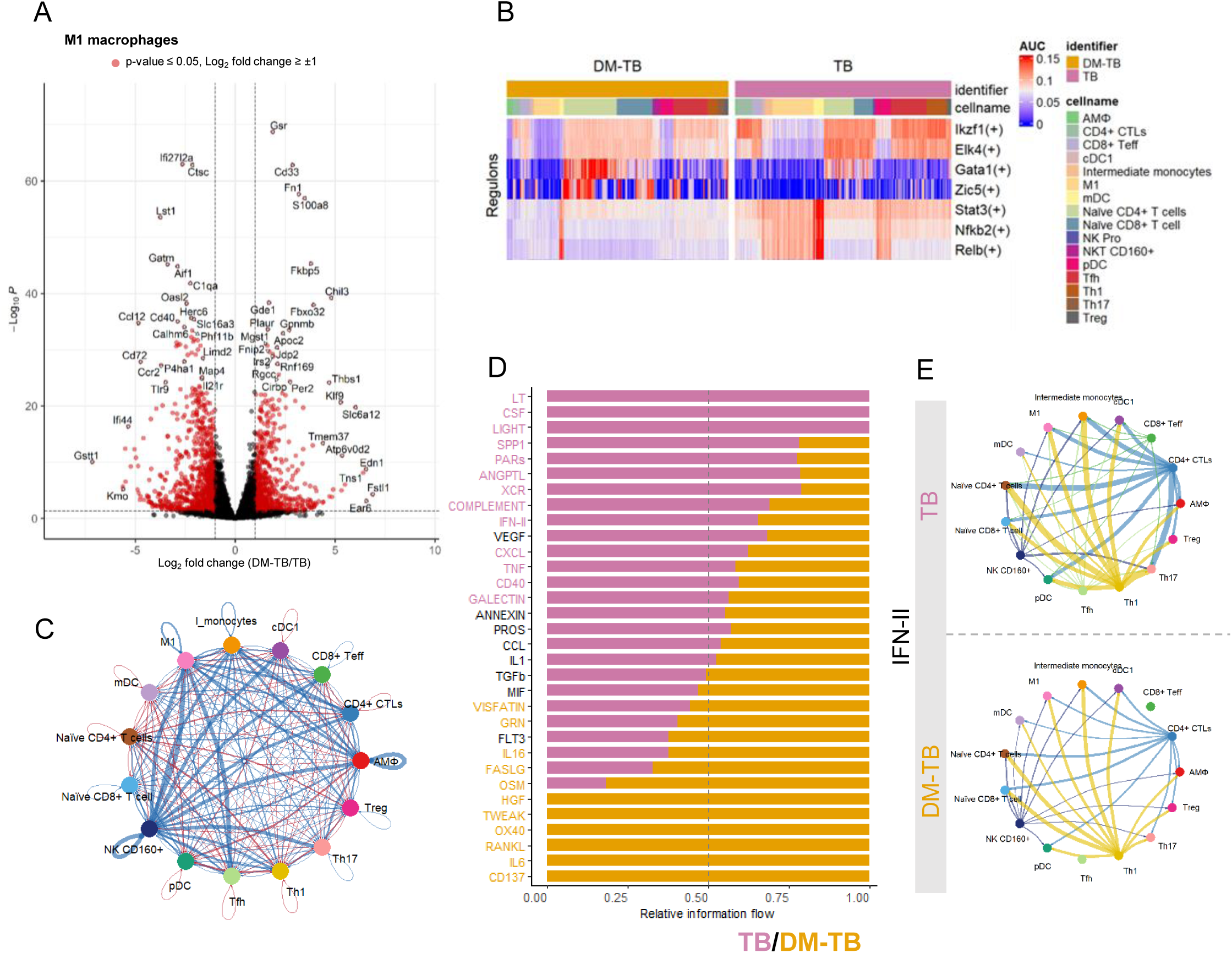
Impaired anti-tuberculosis responses in diabetic mice following Mtb H37Rv infection. (A) Volcano plot showing significantly up-/down-regulated genes in M1 macrophages between DM-TB and TB mice. Significantly deregulated genes: p-value ≤ 0.05, log_2_fold change ≥ ± 1. (B) Heat map showing a subset differentially activated regulon for cell clusters in the DM-TB and TB groups. An unpaired Wilcoxon rank-sum test was performed to compare the activity of each regulon between datasets (TB and DM-TB). Significant difference in regulon activity: p-value ≤ 0.05. (C) CellChat circle plot showing signaling networks in DM-TB and TB groups. The thickness of the edge represents the strength of signaling, and the red and blue colours represent upregulated and downregulated interaction, respectively, in the DM-TB group compared to the TB control. (D) A stacked bar plot showing relative information flow between ligand-receptor pairs in the DM-TB and TB groups. Paired Wilcoxon test was performed to determine whether there was a significant difference in the signaling information flow between healthy and DM groups. The top signaling pathways coloured yellow are enriched in DM-TB, and those coloured pink are enriched in the TB group. Significant difference in information flow: p-value ≤ 0.05. (E) CellChat circle plot showing IFN-II signaling network in DM-TB and TB groups. The thickness of the edge represents the strength of signaling.

The CD4+ CTLs from the lung of the DM-TB group showed 728 DEGs (324/404: down/up, Supplementary Figure S10, Supplementary Table S11). CD4+ CTLs from the lung of the DM group showed activated T-cell homeostatic processes, negative regulation of wound healing and regulation of T-cell apoptotic processes. In addition, pathways like nitric oxide-mediated signaling, glycoprotein metabolic process, and response to copper and zinc ions were also positively enriched. In contrast, processes like pyroptosis, defence response to bacteria, complement activation, and response to IL-1β and IFN-β were suppressed in CD4+ CTLs for the DM-TB group (Supplementary Figure S10, Supplementary Table S12). A suppressed expression of genes involved in complement activation, pyroptosis, leukocyte degranulation and neutrophil activation in CD4+ CTLs points towards a compromised immune response in DM-TB condition.

### Transcriptional regulation of lung immune cells in diabetes and tuberculosis comorbid condition

The Ikzf1, Elk4 regulon activity was suppressed in the lung immune cells of the DM-TB and TB groups (Figure 6B, Supplementary Figure S11, Supplementary Table S13). In contrast, Gata1 and Zic5 regulons were activated in naïve CD8+ and naïve CD4+ cells of DM-TB group (Figure 6B). The Ikzf1 gene is translated to protein Ikaros, which plays a crucial role in T-cell differentiation and early T-cell activation, while Elk4 is important for T-cell activation and immune response upon TCR engagement (Georgopoulos, Winandy, and Avitahl, 1997; Heizmann, Kastner, and Chan, 2018; Yordy and Muise-Helmericks, 2000). Downregulation of these transcriptional pathways could indicate dysfunctional T-cell differentiation and activation in DM-TB conditions. Stat3 is required for T-cell differentiation into Th17 and Tfh cells. Stat3 also has an anti-apoptotic effect on T cells and aids T-cell proliferation and memory formation. In low glucose environments, STAT3 is activated, which promotes FAO, whereas in high glucose environments, STAT3 is inhibited.(Kaminskiy and Melenhorst, 2022) We observed suppression of Stat3 activity across all cell types identified in DM-TB (Figure 6B). Nfkb2 and non-canonical NF-κB signalling mediator Relb activities were downregulated in myeloid cells (AM , cDc1, intermediate monocytes, M1 macrophages, mDCs and pDCs) of the DM-TB group (Figure 6B). NF-κB2 and its’ downstream regulator Relb facilitate activation of antigen-specific T-cells. Relb helps dendritic cells activate T-cells via antigen presentation and non-conventional cross-priming pathway (Bonizzi and Karin, 2004; Sun, 2017). Elevated Relb expression is also correlated with DC maturation. Atf1 and Ctcf regulon activity was also suppressed in all T cell subtypes (CD4+ CTLs, naïve CD4+ T cells, naïve CD8+ T cells, Tfh, Th1, Th17 and Treg cells) of the DM-TB group (Figure 6B). The consistent downregulation of these transcription factors in the DM-TB group indicates a defective functional state of lung immune cells with impaired DC maturity, antigen presentation, impaired T-cell survival, differentiation and a loss of quiescence in naïve T-cells.

### Disrupted cell-cell interactions in diabetes and tuberculosis comorbid condition

Compared to uninfected controls (DM and healthy), both DM-TB and TB groups displayed an increase in overall cell-cell interaction (Figure 6C, Supplementary Figure S5, S12). However, the DM-TB group had more cell-cell communication interactions and altered strength of interactions than the TB group (Supplementary Figure S11). In DM-TB group, signaling by myeloid DCs, classical DCs, CD4+ CTLs, Th1, Tfh, plasmacytoid DCs, and naïve CD4+ T cells was increased. In contrast, signals sent by M1 macrophages, intermediate monocytes, CD8+ effector T cells, alveolar macrophages, regulatory T-cells, Th17 cells, NK CD160+ cells and naïve CD8 T-cells were lower as compared to the TB group (Supplementary Figure S12).

Comparing the overall information flow of each signaling pathway for DM-TB and TB groups revealed increased IL-6, IL-16, TWEAK, RANKL, OX40 and CD137 in DM-TB group (Figure 6D, Supplementary Table S14). In contrast, TNF, CD40, IFN-II, COMPLEMENT, CXCL, and LIGHT signaling were significantly higher in the TB group (Figure 6D). Upon exploring the outgoing signals individually for DM-TB and TB groups, we observed that TWEAK signaling originated mainly from M1 macrophages, OX40 and IL-6 signaling was mediated by mDCs and RANKL was mediated by Tfh and Th17 cells, while all these signals were absent in TB group (Supplementary Figure S12). Mtb hijacks host IL-16 to facilitate its’ intracellular growth (Su et al., 2024). While IL-16 signaling was observed in both DM-TB and TB groups, there was a significant increase in IL-16 mediated signaling for the DM-TB group (Supplementary Figure S11, S12). While Il-1, MIf and TGF-β signaling was similarly active in both groups, IFN-II, TNF and complement signaling, key defenses against Mtb, were significantly reduced in DM-TB group. Exploring the outgoing signal pattern showed that Th1 cells and CD4+ CTLs were majorly responsible for IFN-II signalling in the TB group, supported by NK CD160+ cells, CD8+ effector and naïve CD8+ T cells (Figure 6E). Meanwhile, alveolar macrophages, CD4+ CTLs and Th17 cells were major of TNF signaling (Supplementary Figure S12). CellChat circle plots showed a significantly reduced TNF and IFN-II signaling strength in the DM-TB compared to the TB group (Figure 6E, Supplementary Figure S12). Reduced IFN-II, TNF, and complement signaling, alongside increased IL-6 and IL-16 communication, indicate a skewed inflammatory response in the DM-TB group, that may be detrimental to the host.

### Chronic TB infection in DM condition leads to increased bacterial burden

To investigate the consequences of long-term chronic TB infection in DM mice, we repeated the entire experiment with an additional 56 d.p.i. time point (Figure 7A). In a recent study, Martinez et al. showed that diabetic mice developed using a high-fat diet (HFD) model showed significantly increased lung CFU at 8 w.p.i. (56 d.p.i.) with Mtb Erdman.(Martinez et al., 2023) In the repeat experiment, STZ-NA induced diabetic mice showed an expected reduction in body weight with consistent hyperglycemia over the study period (Figure 7B, 7C). At 21 d.p.i., we did not observe a significant lung or spleen weight. The lung mycobacterial burden was similar between TB and DM-TB groups at 21 d.p.i. corroborating earlier findings (Figure 7D, Figure 7E). However, at 56 d.p.i., mild splenomegaly and a significant increase in the lung weight of DM-TB mice was observed, which indicates a hyper-inflammatory state (Figure 7F), also evident in H&E-stained lung tissue sections for this group (Figure 7H, 7I). At 56 d.p.i., an increased lung mycobacterial load in DM-TB mice was observed, corroborating earlier reports (Figure 7G) (Martinez and Kornfeld, 2014; Martinez et al., 2023).

**Figure 7:**
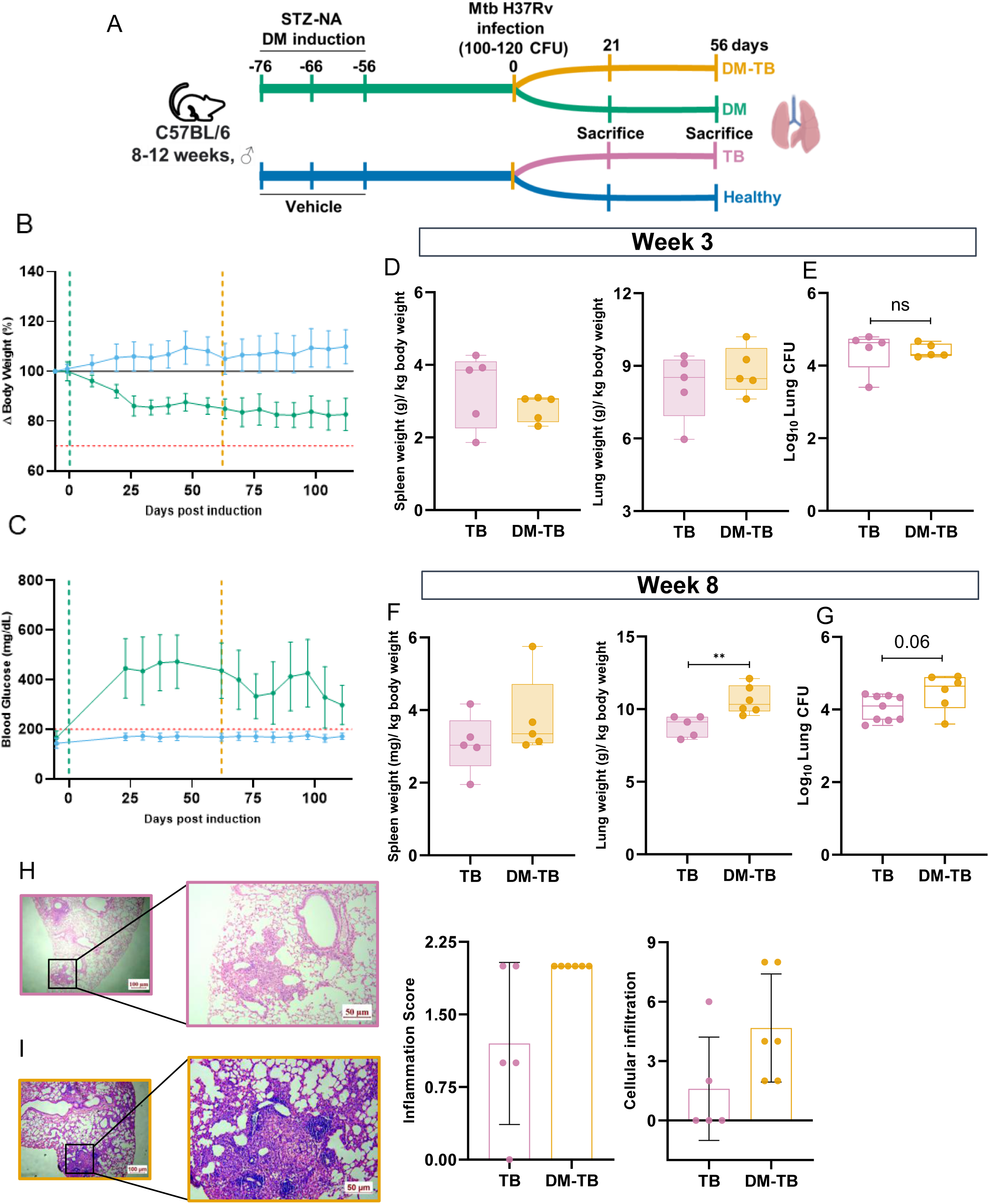
Chronic Mtb H37Rv infection in diabetic mice increases bacterial burden and worsens tissue pathology. A) Schematic representation of the experimental plan. (B) Body weight change in DM and healthy mice during the course of the experiment. Healthy, n = 20; DM, n= 11. Error bars represent standard deviation. (C) Fasting blood glucose levels in diabetic mellitus (DM) mice and healthy controls during the experiment. Healthy, n = 15; DM, n= 21. Error bars represent standard deviation. (D) Spleen and lung weight in TB and DM-TB groups at 3 weeks post-infection. TB, n = 5; DM-TB, n= 5. Error bars represent standard deviation. (E) Lung mycobacterial burden in TB and DM-TB groups at 3 weeks post-infection. TB, n = 5; DM-TB, n= 5. Error bars represent standard deviation. (F) Spleen and lung weight in TB and DM-TB groups at 8 weeks post-infection. TB, n = 5; DM-TB, n= 6. Error bars represent standard deviation. (G) Lung mycobacterial burden in TB and DM-TB groups at 8 weeks post-infection. TB, n = 9; DM-TB, n= 6. Error bars represent standard deviation. (H, I) Hematoxylin and Eosin-stained lung tissue sections with inflammation and cellular infiltration scores of TB and DM-TB groups at 8 weeks post-infection. Statistics: (D-I) unpaired t-test; **: p < 0.005., ns: p > 0.05 at 95% confidence.

## Discussion

Hyperglycemia is a characteristic feature of T2DM and has been associated with microvascular and macrovascular complications such as atherosclerosis, retinopathy, nephropathy and neuropathy (DeFronzo et al., 2015; E. Dale Abel, 2024; Katsarou et al., 2017; Papatheodorou et al., 2018). Diabetes also renders the host susceptible to several pathogenic attacks like COVID-19, candida, influenza, Mtb and *Helicobacter pylori* (Berbudi et al., 2020; Daryabor et al., 2020; Papatheodorou et al., 2018). This increased susceptibility to infections is owed to disrupted immune responses in diabetic conditions, such as impaired immune cell functions like chemotaxis, reduced phagocytic ability and insufficient cytokine response (Berbudi et al., 2020; Daryabor et al., 2020; Papatheodorou et al., 2018).

To investigate how diabetes influences lung immune dynamics, we used the STZ-NA model, which induces T2DM-like hyperglycemia (Cheekatla et al., 2016). Unlike genetic and diet-based models that take longer to develop hyperglycemia and often lead to obesity, STZ-NA–treated mice become hyperglycemic within 72 hours and remain so, exhibiting T2DM-associated complications (Martinez et al., 2023). In addition, both genetic and diet-based models of diabetes lead to obesity, which contributes significantly towards organ dysfunction and systemic inflammation. We aimed to understand the effects of hyperglycemia on host metabolism and immune dysfunction in the absence of confounding factors contributed by obesity. The DM induction method adopted in this study was a modified version of STZ-NA T2DM induction reported earlier (Cheekatla et al., 2016). Since Cheekatla et al. reported resistance to DM development in female mice, in this study, male mice were used with a reduced STZ dose of 150 mg/kg (Cheekatla et al., 2016). Male C57BL/6 and BALB/c mice are reported to have a higher susceptibility to H37Rv infection with increased morbidity and mortality; thus, male mice were used in this study (Hertz et al., 2020). Over one month of monitoring, these mice maintained persistent hyperglycemia, accompanied by significant insulin resistance, glucose intolerance, and dyslipidemia compared to controls. While female C57BL/6 mice develop similar metabolic disruptions over six months (Cheekatla et al., 2016)), our male mice displayed them within a month, underscoring the role of sex in faster, more severe disease progression.

Following low dose (100-120 CFU) Mtb aerosol infection, DM-TB and TB groups showed similar tissue mycobacterial burden at 3 w.p.i., corroborating earlier reports (Cheekatla et al., 2016). The gross tissue pathology and the spleen weight were similar between the DM-TB and TB groups. Upon histopathology analysis, both groups exhibited cellular infiltration, but granulomatous structures were absent in either group. However, upon chronic infection, i.e., 8 w.p.i., DM-TB mice developed increased lung mycobacterial burden with significantly increased lung weight, indicating hyperinflammation.

We observed a significantly perturbed serum cytokine profile in diabetic and DM-TB conditions, corroborating earlier findings (Domingo-Gonzalez et al., 2016). Upon Mtb infection, circulatory IFN-γ levels significantly increased in DM-TB and TB groups compared to the uninfected controls. However, this increase was lower in the DM-TB group than in TB controls, suggesting a dampened IFN-γ mediated anti-TB response. Circulatory TNF-α was also significantly higher in DM, TB and DM-TB groups compared to the healthy controls with highest levels in the DM-TB group, likely due to infection-mediated augmentation compounding preexisting TNF-α levels due to diabetes. While TNF-α is required by the host to combat Mtb infection, excess of it has been shown to increase susceptibility to the pathogen. Therefore, the increase in circulatory TNF-α levels due to cumulative effects of T2DM and TB could prove detrimental to the host due to higher susceptibility to pathogens and worsening insulin-resistant state (Roca et al., 2022; Stephens, Lee, and Pilch, 1997).

Furthermore, circulatory IL-6 levels were significantly increased in DM and DM-TB groups than in healthy controls echoing earlier reports linking IL-6 to impaired insulin signaling in diabetes (Fasshauer and Paschke, 2003; Kim et al., 2008). IL-6 is also involved in early-stage immune responses in Mtb infections and is responsible for accumulating lymphocytes and regulating mononuclear cell-driven inflammation (Li, Jones, and Geiger, 2018). However, increased baseline IL-6 levels in DM-TB conditions have been associated with worsened treatment outcomes (Gupte et al., 2022). In addition, we also observed significantly increased circulatory IL-17A levels in DM and DM-TB groups, potentially driving chronic inflammation in diabetes through NF-κB activation (Abdel-Moneim, Bakery, and Allam, 2018; Merino, Jazwinski, and Rout, 2021). A modest increase in circulatory IL-1β levels in the DM-TB group compared to controls further supports the presence of an underlying chronic inflammatory state. Altogether, these data suggest that elevated IL-6, TNF-α and IL-17A in STZ-NA induced DM and DM-TB conditions could be drivers of tissue pathology, exacerbated glycemic control and enhanced insulin resistance.

The dissemination of Mtb precedes T-cell immunity (Chackerian et al., 2002). Upon infection, Mtb enters host immune cells like macrophages and dendritic cells and is transported to pulmonary lymph nodes (PLN),where T-cells are activated and migrate to the lungs (Chackerian et al., 2002). Thus, macrophages, DCs and T-cells are central to the primary immune response against Mtb. To investigate these responses, we sorted lung CD11c+ (macrophages and DCs) and CD3+ (T cells) cells at 21 days post-infection (3 w.p.i.) from the lungs of euglycemic (TB) and hyperglycemic groups (DM-TB, DM) and control mice (healthy).

Consistent with earlier findings, diabetes delayed T-cell migration to the site of infection (Vallerskog, Martens, and Kornfeld, 2010). While both CD11c+ and CD3+ cells were significantly increased with Mtb infection, DM-TB group showed significantly reduced CD3+ cell frequency than TB group, indicating delayed T-cell recruitment. In contrast, CD11c+ cell frequency in the DM-TB group was lower than TB, although statistically insignificant. An overall increase in the proportion of naïve CD4+ T-cells and Th17 cells in the DM group was observed. Increased frequency of Th17 cells has been associated with inflammageing, defined as chronic low-grade inflammation in the absence of infection accompanying ageing. T2DM has been associated with the early onset of inflammageing, even in the absence of obesity (Prattichizzo et al., 2018). Reports show that aged individuals and T2DM patients present with increased Th17 frequencies (Okamoto Yoshida et al., 2010). Increased Th17 cell numbers are accompanied by higher circulatory IL-17A and IL-6 levels, and our findings corroborate earlier literature. IL-17 knockout mice models demonstrate improved insulin sensitivity, glucose tolerance and reduced pro-inflammatory cytokine levels (Abdel-Moneim, Bakery, and Allam, 2018; Okamoto Yoshida et al., 2010). Transplanting islet-reactive Th17 cells is known to accelerate the onset of diabetes in non-obese diabetic (NOD) mice (Abdel-Moneim, Bakery, and Allam, 2018). Although, most studies have profiled circulatory immune cells, we report that T2DM leads to increased Th17 cell number in the lungs of diabetic mice, contributing to low-grade chronic inflammation.

We observed that genes associated with pyroptosis were significantly upregulated in naïve CD4+ T-cells from the lungs of DM mice compared to healthy controls. Pyroptosis, a gasdermin mediated cell death mechanism, has been implicated in various sterile inflammatory disorders such as T2DM, cancer, atherosclerosis and lung inflammation (Li et al., 2021). Although inflammasome-mediated pyroptosis of pancreatic β-cells is well-known in T2DM (Li et al., 2021), our findings indicate that pyroptosis-mediated inflammation in T2DM is also prevalent in the lungs and not just confined to the pancreas. In our earlier work, we highlighted the multi-tissue metabolomic dysregulation in DM-TB conditions, and it might be interesting to see the multi-tissue immunological consequences of T2DM (Chaudhary, Pahwa, and Nanda, 2024).

Th17 cells, intermediate monocytes and naïve CD8+ T cells emerged as major drivers of increased IL-16 signalling in the lungs of diabetic mice. In ova-induced mice models of asthma, IL-16 has been identified as a major driver of airway inflammation and its increased signaling in immune cells from diabetic mice suggests increased baseline respiratory inflammation (Li et al., 2019a). Supporting this observation, histology analysis of lung tissue sections from 3 and 8 w.p.i. mice exhibited increased tissue inflammation in DM-TB conditions, where the increased IL-16 signalling could be the major driver and warrants further investigation.

Naïve CD4+ T-cells and Th17 cells were more abundant in the lungs of DM-TB mice than in TB alone, while the number of M1 macrophages, Th1 cells and CD4+ CTLs were lower. Th1 cells secrete IFN-γ, which activates macrophages to mediate Mtb killing and also play an anti-inflammatory role by restricting tissue-damaging IL-17 responses (Lyadova and Panteleev, 2015).

Increased CD4+ CTLs presence in the circulation of drug-sensitive active TB patients was reported recently (Flores-Gonzalez et al., 2024). While these cells have been studied extensively in viral infections (dengue, influenza, hepatitis and HIV), which exert cytotoxic and protective roles, their role in bacterial infections, particularly Mtb, remains underexplored (Cenerenti et al., 2022). CD4+ CTLs, like Th1 cells, produce IFN-γ secreting cells and might be involved in anti-TB activity by activating macrophages. In addition, these cells act majorly via MHC-II interactions and, therefore, might be involved in the killing of Mtb harbouring macrophages in the host lungs via the secretion of cytolytic molecules such as granzyme (Cenerenti et al., 2022).

M1 macrophages play a crucial role in pro-inflammatory host responses against intracellular bacteria. These cells exhibit high iNOS expression, leading to NO production, which is detrimental to Mtb and when polarised by IFN-γ, secrete pro-inflammatory cytokines, anti-microbial peptides and proteolytic enzymes (Khan et al., 2022; Lugo-Villarino et al., 2011). However, Mtb blocks M1 polarisation by restricting the expression of IFN-γ responsive genes by inducing secretion of IL-6 (Boni et al., 2022). Mtb ESAT-6 also limits NF-κB2 and IFN-γ regulatory factors downstream of TLR2 signalling. In our SCENIC analysis, NF-κB2 activity was significantly downregulated in M1 and alveolar macrophages from the DM-TB group compared to TB controls, while CellChat analysis identified a reduced MIF signalling pathway in M1 M from DM-TB mice. It is worth noting that both TL2 signalling and MIF signalling are regulated via NF-κB2, and compromised activation of NF-κB2points towards a disrupted M1 functionality in DM-TB condition. Overall, lower Th1, M1 M and CD4+ CTLs populations in the lung of DM-TB mice indicate an impaired anti-TB immune response in this group.

In the current study, we have attempted to elucidate the lung immune landscape of DM-TB comorbid conditions. However, the current study has a few limitations. Gender-based differences are reported in the context of metabolic as well as immunological consequences of DM-TB comorbidity. In this study, male C57BL/6 mice were used for the immunological studies. Expanding this work to female mice would greatly enrich the insights gathered from the current work. As the current study used chemical diabetes induction methods, employing alternate DM models such as HFD-induced or genetic models (Akita mice, db/db mice) would help validate these findings. In addition, the effect of anti-diabetic drugs such as insulin, Glucagon-like peptide (GLP) inhibitors or metformin on immunological consequences of DM-TB comorbid condition remains to be explored. The observed deregulation of cytokines (MIF, IL-16 and TGF-β) in DM and their crucial role in host TB defence needs to be validated using monoclonal antibodies to observe if they result in reduced disease severity in DM-TB comorbidity and needs further exploration.

However, our findings reveal that DM-TB comorbidity results in disrupted immunological host profiles compared to euglycemic controls. A dampened immune response to Mtb by the diabetes condition might increase the disease severity in comorbid individuals. This work provides an extensive baseline dataset, offering cellular and molecular insights to enhance our understanding of the DM-TB comorbid condition. Subsequent studies involving human subjects would be crucial to validate these results and instrumental in designing targeted drug interventions.

## Materials and Methods

### Diabetes induction

The animal experimental protocols followed in this study are approved by the animal ethics committee (vide reference ICGEB/IAEC/07032020/TH-13Ext) of the International Centre for Genetic Engineering and Biotechnology New Delhi component. As reported earlier, DM was induced in male C57BL/6 mice of 8-12 weeks of age by combined administration of STZ and NA, with minor modifications (Cheekatla et al., 2016). Mice were fasted for 16 hours before NA and STZ administration. NA (60 mg/kg in 0.9% saline) was administered intraperitoneally 15 minutes before STZ injection. STZ (150 mg/kg in 50mM citric acid buffer) and NA were administered intraperitoneally thrice with an interval of 10 days between doses. Age and gender matched control mice received intraperitoneal injections of the corresponding vehicle solution. Mice with fasting plasma glucose >200 mg/dL (measured using Dr. Morepen Gluco-One BG-03 glucometer) were grouped as diabetic (DM). Body weight and 6 hours of fasting blood glucose were monitored weekly till experiment completion.

### Intra peritoneal Glucose tolerance test (iPGTT) and Insulin Tolerance Test (ITT)

Mice were fasted for 6 hours with access to drinking water at all times. For iPGTT, mice were weighed and intra-peritoneally injected with a 20% glucose solution (2g of glucose/kg body mass). For ITT, mice were weighed, and the volume of 0.25 UI insulin solution required for IP injection was calculated using the formula: volume of IP insulin injection (μl) = 3 x body weight (g). The baseline glucose level (t = 0) was measured using the Dr. Morepen Gluco-One BG-03 glucometer. Mice were injected intraperitoneally with the appropriate amount of glucose solution, as previously determined, and the time-point of injection was recorded. Blood glucose levels were measured 15, 30, 60, 90 and 120 minutes after glucose injection by placing a small drop of blood on a new test strip and recording the measurements. At the end of the experimental session, water and food were provided ad libitum.

### Mtb H37Rv infection

After 60 days of consistent hyperglycemia, C57Bl/6 mice were infected with a low aerosol dose (100-120 CFU/animal) of animal passaged Mtb H37Rv using a Madison chamber in a tuberculosis aerosol challenge BSL-III facility at the host institution. Before infection, Mtb H37Rv was grown to log phase in 7H9 medium supplemented with OADC (10%, Difco Middlebrook). The PDIM levels of the Mtb H37Rv strain were monitored using standard procedure. For infection, bacterial culture (10 ml) was washed with Phosphate Buffer Saline (PBS, pH 7.4, 9 ml) and centrifuged at 4,000 g for 10 minutes at room temperature. After discarding the supernatant, the resulting pellet was resuspended using PBS (1 ml). The bacterial single-cell suspension was prepared in PBS (15 ml) and adjusted to 15 × 10^6^ cells/ml before loading to the nebuliser chamber for infection. Study mice were exposed to aerosol for 20 minutes in a Madison chamber, which delivers 100-120 Colony-forming units (CFU)/animal. After 1-, 21- and 56 days post-infection (d.p.i.; 3- and 8-weeks post infection), animals were anaesthetized for whole blood collection via retro-orbital puncture and serum was isolated. At 21 and 56 d.p.i., organs (lung, liver and spleen) were harvested for bacterial CFU assay, histology and immune cell isolation. Part of the lungs was stored in buffered formalin solution for histology analysis using standard hematoxylin and eosin staining methods. For CFU enumeration, the spleen, lung and liver were homogenized in phosphate buffer saline (PBS) and plated on 7H11 media supplemented with OADC (10%), BBL MGIT PANTA and thiophene carboxylic acid hydrazide (TCH; 20 µg/ml) in triplicates and CFU counting was carried out 21 days later.

### Serum insulin, fatty acids, cholesterol and triglyceride measurements

Mice were anaesthetized using isoflurane and whole blood was collected via retro-orbital puncture for all study animals. Whole blood was allowed to clot at room temperature for 20 minutes and then centrifuged at 2,000 rpm at 4°C for 10 minutes. Serum was transferred to a fresh micro-centrifuge tube and stored at -80°C until further processing. Random serum insulin levels of all mice groups were measured using a Rat / Mouse Insulin ELISA Kit (#EZRMI-13K, Merck Millipore, USA) following the manufacturer’s instructions. Serum-free fatty acids (#700310), cholesterol (#10007640), and triglyceride (#10010303) levels were measured using either a fluorometric or colorimetric assay (Cayman Chemicals, USA), following the manufacturer’s instructions.

### Cytokine measurement and analysis using LEGENDplex^TM^

Test serum samples were thawed on ice and centrifuged at 2,000 rpm for 5 minutes at 4°C. Wash buffer, matrix C and standards were prepared per the manufacturer’s instructions. Serum samples (25 µl) were diluted 2-fold with assay buffer, and cytokine (IFN-γ, Il-6, TNF-α, IL-17A, IL-23, IL-1α and IL-1β) estimation was performed as per manufacturer’s instructions (BioLegend, California, USA).

### Lung immune cell isolation

Lungs of study groups were chopped into smaller pieces using surgical scissors and digested at 37°C and 180 rpm for 30 minutes in RPMI (Gibco, 31800022) containing collagenase D (2 µg/µl, Sigma-Aldrich Chemicals Pvt Limited, 101088866001) and DNase-I (1 µg/µl, Sigma-Aldrich Chemicals Pvt Limited, 10104159001). Following incubation, samples were passed through a 70 μm nylon mesh cell strainer, and the single-cell suspension was collected in a 15 ml centrifuge tube. Samples were centrifuged at 250 g at 4°C for 5 min, and the resulting cell pellet was resuspended in ACK lysis solution (1 ml). Samples were incubated for 3 min at room temperature, and lysis was stopped by adding ice-cold 10% FACS buffer (2 ml) to each sample. Samples were centrifuged at 250 g at 4°C for 5 min, and the pellet was resuspended in 1 mL of 10% FACS buffer. Cells were enumerated using Countess 3 (Thermo Fisher Scientific, USA), and viability was determined by trypan blue staining.

### Cell surface protein staining and FACS sorting

After transferring the cells to a new 2-ml microcentrifuge tube, it was centrifuged at 250 × g at 4°C for 5 min. The pellet was resuspended in 94 μl chilled PBS supplemented with FBS (10%). TruStain FcX (0.5 μl) and True-Stain Monocyte Blocker (5 μl) were added to the cells and incubated for 10 min at 4°C. To respective samples, BioLegend TotalSeq^TM^-B hashtags (0.5 μl) were added and incubated for 30 min at 4°C. Samples were diluted with PBS supplemented with FBS (10%), and after making the final volume to 2 ml were centrifuged at 250 × g at 4°C for 5 min. The pellet was resuspended in diluted live/dead stain (98.5 μl) and FACS antibody mix and incubated for 45 min at 4°C without light exposure. Samples, after diluting in PBS supplemented with FBS (10%) were made to the final volume of 2 ml and centrifuged at 250 g at 4°C for 5 min. The pellet was resuspended in 1.5 ml of chilled PBS supplemented with FBS (10%) and transferred to 5 ml round-bottom polystyrene tubes and flow sorted cells were collected in 1 ml chilled PBS supplemented with FBS (50%).

### Sample Fixation for scRNAseq

Flow sorted cells were counted using Countess 3 (Thermofisher Scientific, USA). CD3+ and CD11C+ cells were mixed in a ratio of 2:1 in a 15 ml centrifuge tube and centrifuged at 250 g at 4°C for 5 min. The supernatant was discarded, and cells were fixed, following the manufacturer’s protocol. Briefly, cells were resuspended in room temperature fixation buffer (1 ml, 10x Genomics PN-2000517) and incubated at 4°C for 24 hours. Samples were then centrifuged at 250 g at 4°C for 5 min, resuspended in quenching solution (1 ml, 10x Genomics, USA PN-2000516) and transferred to a 2 ml DNA LoBind (022431048, Eppendorf) tube. Enhancer (10x Genomics, PN-2000482) was thawed and pre-warmed at 65°C for 10 minutes. Glycerol (50%, 275 µl, G5516, Millipore Sigma) and enhancer (100 µl) were added to the quenched cells and samples were stored at -80°C until further processing.

### Fixed single-cell RNA gene expression profiling

The fixed cells stored at -80°C were used for fixed single-cell RNA sequencing 10x Genomics single-cell gene expression flex kit. Probe hybridization, GEM generation, barcoding and library construction were performed following manufacturer’s instructions. Sample libraries were stored at -20°C until sequencing, and an aliquot (1 μl) was used for library QC using D1000 ScreenTape with Agilent TapeStation (Agilent Technologies, USA). Gene expression (GEX) and cell surface protein (CSP) libraries were sequenced using PE150 chemistry on an SP100 flow cell in a third-party NovaSeq 6000 sequencing system (Illumina, USA). The sequencing depth was 10,000 read pairs/cell for GEX libraries and 5,000 read pairs/cell for CSP libraries. A 10% PhiX spike was used while loading for the sequencing of these libraries.

### scRNAseq data analysis

All the scRNA-seq raw and processed data files are accessible from Gene Expression Omnibus (GEO) database with accession number GSE280089. FASTQ files obtained from the NovaSeq6000 sequencing system were run through Cell Ranger (v7.2) multi pipeline. After successfully running the Cell Ranger multi pipeline, the output folder “filtered_feature_bc_matrix” generated for each sample was used for further data analysis using the Seurat V5 package in R. The scRNA seq data analysis was divided majorly into Mtb infected and uninfected control groups. Samples were read using the Read10X function of Seurat. Data was subsetted using the following filter parameters: nFeature_RNA (no. of features/cell) = 1000-7000, nCount_RNA (UMI count/cell) =1500-20000 and percent.mt (mitochondrial RNA percentage) ≤ 5. Data was normalised using the NormalizeData function of the Seurat package, where the count data in RNA assay is log normalised, and a scale factor of 10,000 is used for cell-level normalisation. FindVariableFeatures function was used to identify features in the dataset that were classified as outliers on a ’mean variability plot’. VST (variance stabilising transformation) was used as a selection method, accounting for the variance-mean gene expression relationship with a regression model. For comparative analysis of two different datasets, the Seurat object for each dataset needs to be integrated. Before integrating Seurat objects, the FindIntegrationAnchors function is used to find a set of anchors between a list of Seurat objects. These anchors are then used to integrate the objects using the IntegrateData function. A list of genes to keep in the merged object post-integration was defined to ensure that genes are not subsetted into different layers of the Seurat object post-integration. Then, the IntegrateData function was used to perform dataset integration. features.to.integrate into the IntegrateData was defined as the list of genes prepared in the previous step. DefaultAssay was set to the “integrated” layer of the merged Seurat object for further data analysis. The features in the dataset were scaled and centered using the ScaleData function, where the maximum value to return for scaled data was set at 10. Dimensionality reduction in the dataset was performed using the RunUMAP function, which runs the Uniform Manifold Approximation and Projection (UMAP) dimensionality reduction technique. FindNeighbors function was used to compute the k.param nearest neighbours for the integrated dataset. FindClusters function was used to identify cell clusters by a shared nearest neighbour (SNN) modularity optimization-based clustering algorithm. The custom function was used to plot integrated clusters as bar plots with cluster identity on the X-axis and the number of cells per cluster on the Y-axis. ScType package was used for cluster annotation, a cell-type identification algorithm that uses a comprehensive cell marker database as background information to assign an identity to a cell. The datasets were subsetted into myeloid (Itgam+ Itgax+ Cd13+ Cd14+), and lymphoid (Cd3e+ Cd4+ Cd8a+ Cd8b+) clusters based on canonical marker expression and reclustered to identify further sub-clusters in these cell populations.

### Differential gene expression (DGE) analysis

DESeq2 package was used for differential expression analysis to identify differences in the gene expression between study groups. Before proceeding with DESeq2 analysis, the scRNA seq samples must be pseudobulked. A pseudobulk sample is formed by aggregating the expression values for a group of cells from the biological replicate. Since we used cell surface protein labels for multiplexing four biological replicates for each study group, we fetched the antibody data from cell ranger output and assigned a replicate identity to each cell and the group identity. This was done using custom functions and resulted in merging the replicate information to the metadata of the Seurat object. AggregateExpression function from the Seurat package was then used to pseudobulk data, which returns summed counts for each study group. DGE analysis was performed using the DESeq2 package, which estimates dispersion and fits a negative binomial model to the count data. During analysis, euglycemic control (TB or healthy) groups were set as reference levels to use as a baseline for comparison. DGE analysis was performed for individual cell clusters between study groups to identify diabetes-associated changes in gene expression for each cell type.

### CellChat analysis

CellChat is an R package that uses gene expression data of a cell and models the probability of cell-cell communication.(Jin et al., 2021) This package integrates gene expression with the CellChat database, which details interactions between signaling ligands, receptors and their cofactors. CellChat analysis used cell annotation to identify the study group, and cell-cell communication was computed for each study group independently. CellChatDB.mouse was a database comprising a list of ligand-receptor interactions in the mice. Cell-cell communication probability was computed between interacting cell clusters using the computeCommunProb function with trimean to compute average gene expression per cell group. computeCommunProbPathway function was used to calculate communication probability for a signaling pathway by summarising all related ligands/receptors, and the aggregateNet function was used to calculate aggregated cell-cell communication. Each study group’s resulting CellChat object was exported as a .rds file. The cellchat .rds file for each study group was imported for comparative CellChat analysis, and network centrality was computed using the netAnalysis_computeCentrality function for each sample. The CellChat objects for each study group were merged using the mergeCellChat function. Using the compareInteractions function, the total number of interactions and strength was compared between datasets to assess whether cell-cell communication was altered between groups. netVisual_diffInteraction function was used to differential cell-cell communication between study groups. The ranked function compared the information flow between groups’ signaling pathways.

### SCENIC analysis

The Single-Cell Regulatory Network Inference and Clustering (SCENIC) package allows differential activation states of transcription factors and gene regulatory networks in cell types from scRNA-seq data.(Aibar et al., 2017) SCENIC analysis was done using Python. Outputs from the SCENIC analysis were exported as .loom files. QC-filtered Seurat objects for each dataset were exported as .rds files, and .loom files were prepared from .rds objects and used for SCENIC analysis. Gene regulatory inference was performed using the GRNBoost2 algorithm. This step identifies potential targets for each transcription factor based on co-expression. Next, regulon prediction was performed based on DNA motif analysis. For this, ranking and motif annotations databases for mice were downloaded from the cisTarget resources website. Network activity in individual cells was analysed using AUCell, which allows the identification of cells with active gene sets. The resulting files were used in R for comparative regulon analysis between groups. Integrated .rds object and .csv file containing auc information were imported for study groups, and the unpaired Wilcoxon rank-sum test was performed on each common regulon from study groups to identify significantly deregulated regulons. Combinations with p-value ≤ 0.05 were defined as significant differential regulons and used to plot data as a cluster map.

### Statistical Analysis

Animal body weight, blood glucose, Mtb H37Rv CFU, iPGTT, ITT, cytokine concentration and immune cell frequency data were analysed by parametric analysis using Student’s t-test and area under curve (AUC) analysis in GraphPad Prism (version 8.4.2, GraphPad Software, San Diego, CA). R packages pheatmap, Dimplot, Nebulosa, DotPlot, ggplot2, EnhancedVolcano, ComplexHeatmap, Rcolorbrewer, scCustomize, vlnplot and corrplot were used to generate plots. FlowJo (version 10.7.1, BD Biosciences, Ashland, USA) was used for FACS data analysis. LEGENDplex data was analysed using BioLegend’s LEGENDplex™ data analysis software (Version 2024-06-15, LEGENDplex Qognit).

## Supporting information

Supplementary Figures Revised

## Acknowledgement

SC is a Shyama Prasad Mukherjee fellow and was supported by the Council of Scientific and Industrial Research, Government of India. A fellowship from the Department of Biotechnology, Government of India, supported MS and FP. The core support from the International Centre for Genetic Engineering and Biotechnology (ICGEB), New Delhi, to RKN; core support from the National Institute of Immunology, New Delhi, to DM; core support from the Institute of Life Sciences Bhubaneswar to DD are highly acknowledged. Nidhi Yadav and Ashish Gupta are acknowledged for their help and support during the animal experiments. Help from the staff of the bio-experimentation facility and Tuberculosis Aerosol Challenge Facility (TACF) at ICGEB, New Delhi, is acknowledged. TACF is supported by the Department of Biotechnology, Government of India. We thank Dr. Veena Patil and Mr. Raunak Kar from the National Institute of Immunology, New Delhi, for their help with the single-cell RNA sequencing experiments. For the SCENIC analysis, CSIR-IGIB Tejas HPC compute is acknowledged. Part of these research findings was presented as a poster at the Global Immunology Summit 2024 at THSTI on 22nd February 2024. Some figures have been created with Biorender.com.

## Author Contribution

S. C. and R. N. conceptualization, S. C., M. S., F. P. and R. N. methodology, S. C. visualization, S. C., P.S. and S.V. formal analysis, S. C. and R. N. writing–original draft; S. C., D. M., D. D., R. N. review – final draft, D. M., D. D. and R. N. supervision.

## Data Availability

The single-cell RNA sequencing data generated in this study are available in the Gene Expression Omnibus (GEO) under accession number GSE280089.

## Code Availability

All R scripts used for data analysis and figure generation in this manuscript are accessible on GitHub at https://github.com/ChaudharyShweta/DM-TB-scRNA-seq.

