## Supplementary Figures Revised for "Single-cell profiling of the lung immune cells of diabetes-tuberculosis comorbidity reveals reduced type-II interferon and elevated Th17 responses"

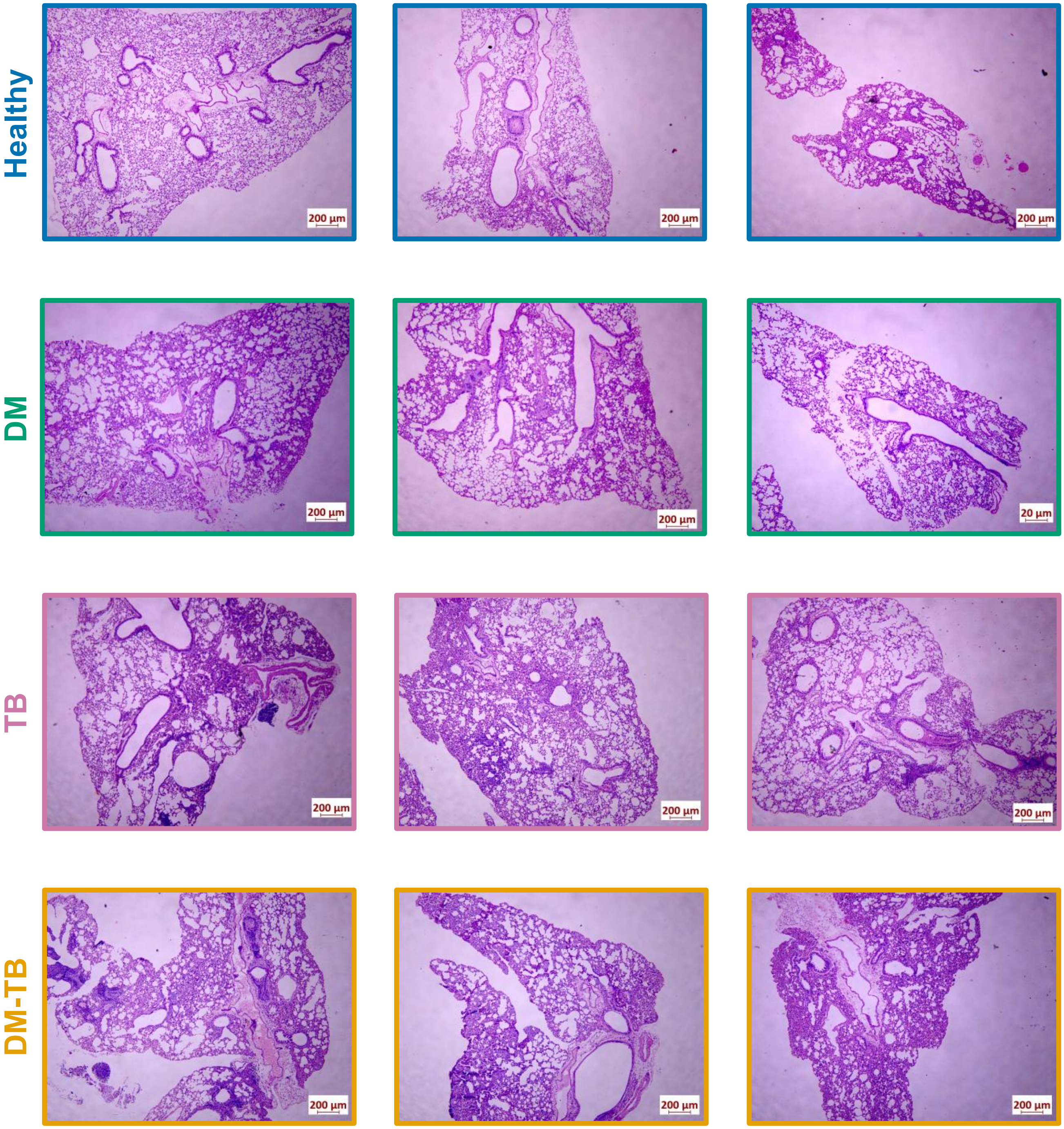


Supplementary Figure S1 continued

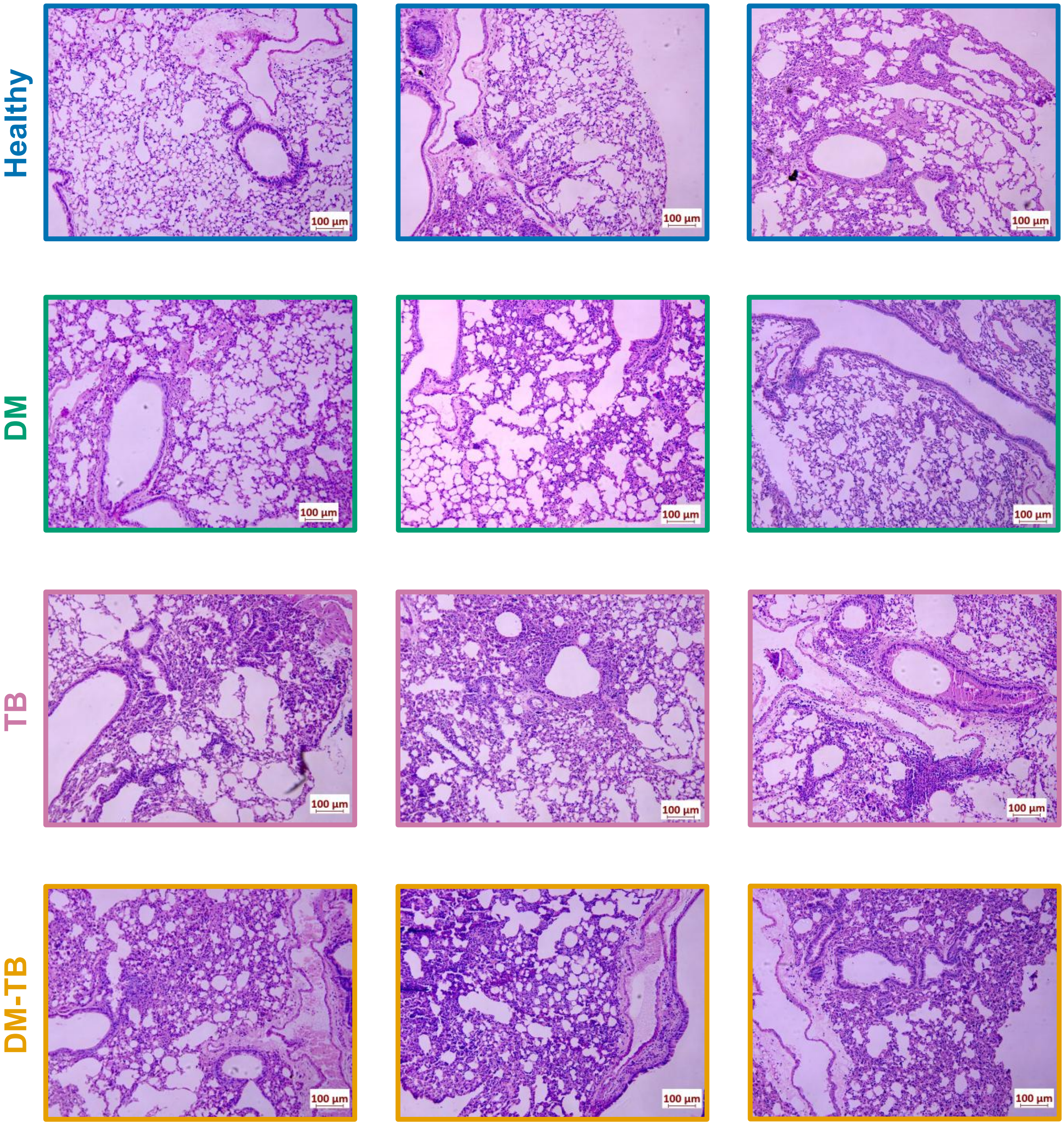


**Supplementary Figure S1:** Hematoxylin and Eosin-stained lung tissue sections showing histopathology differences (inflammation, neutrophil infiltration) between study groups (Healthy, DM: Nicotinamide-Streptozotocin induced diabetic C57BL/6 mice, TB: Mycobacterium tuberculosis H37Rv infected control mice, DM-TB: Mycobacterium tuberculosis H37Rv infected DM mice). Scalebar-lungs:100μm; 200μm.

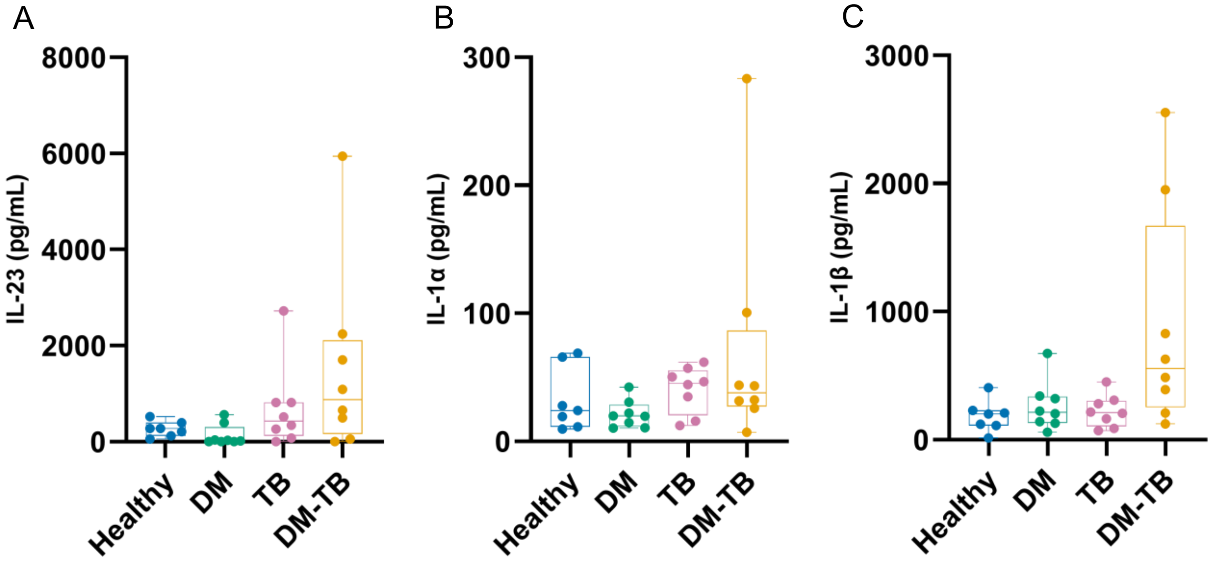


**Supplementary Figure S2:** Serum (A) IL-23, (B) IL-1 α and (C) IL-1β levels (pg/mL) measured via legendplex assay in healthy, DM, TB and DM-TB groups at 21 d.p.i. DM: Nicotinamide-Streptozotocin induced diabetic C57BL/6 mice, TB: Mycobacterium tuberculosis H37Rv infected control mice, DM-TB: Mycobacterium tuberculosis H37Rv infected DM mice

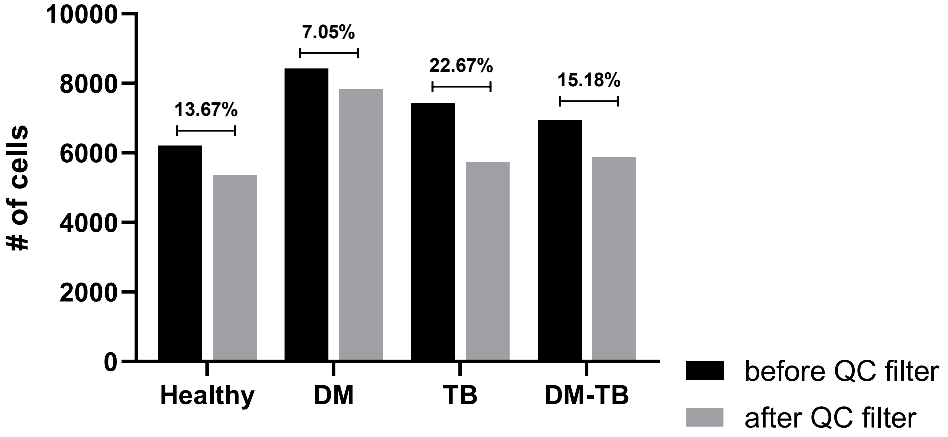


**Supplementary Figure S3:** Percent of flow sorted CD3+ and CD11c+ cells lost after QC filtering and number of cells retained per group. DM: Nicotinamide-Streptozotocin induced diabetic C57BL/6 mice, TB: Mycobacterium tuberculosis H37Rv infected control mice, DM-TB: Mycobacterium tuberculosis H37Rv infected DM mice

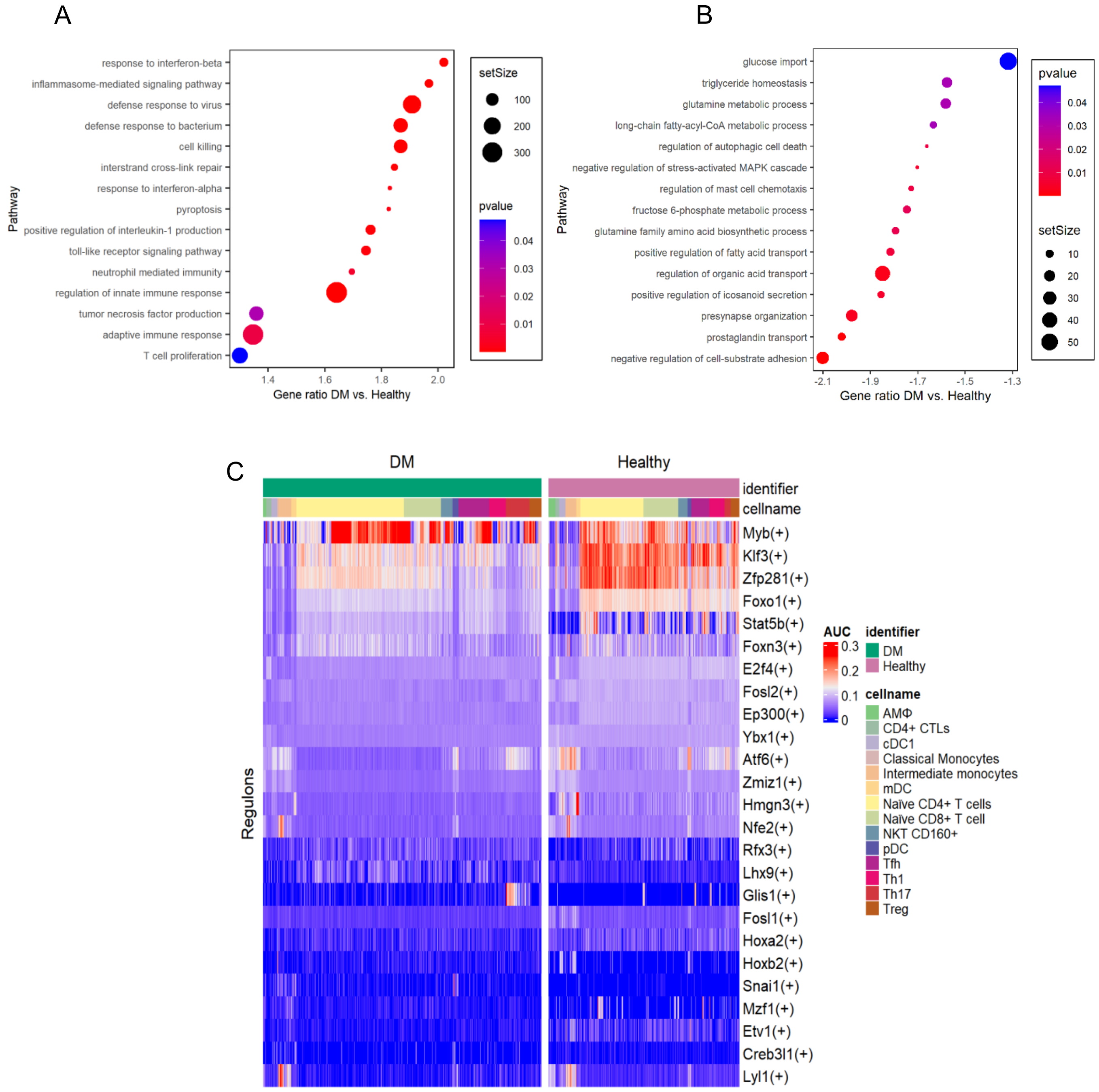


**Supplementary Figure S4:** Gene Set Enrichment Analysis (GSEA) for biological processes of naïve CD4+ T-cells for (A) positively and (B) negatively enriched terms in DM as compared to healthy group (C) Heat map showing differentially activated regulon for cell clusters in hyperglycemic (DM) and euglycemic (healthy) groups. Unpaired Wilcoxon rank-sum test was performed to compare the activity of each regulon between datasets (healthy and DM). Significant difference in regulon activity at p-value ≤ 0.05.

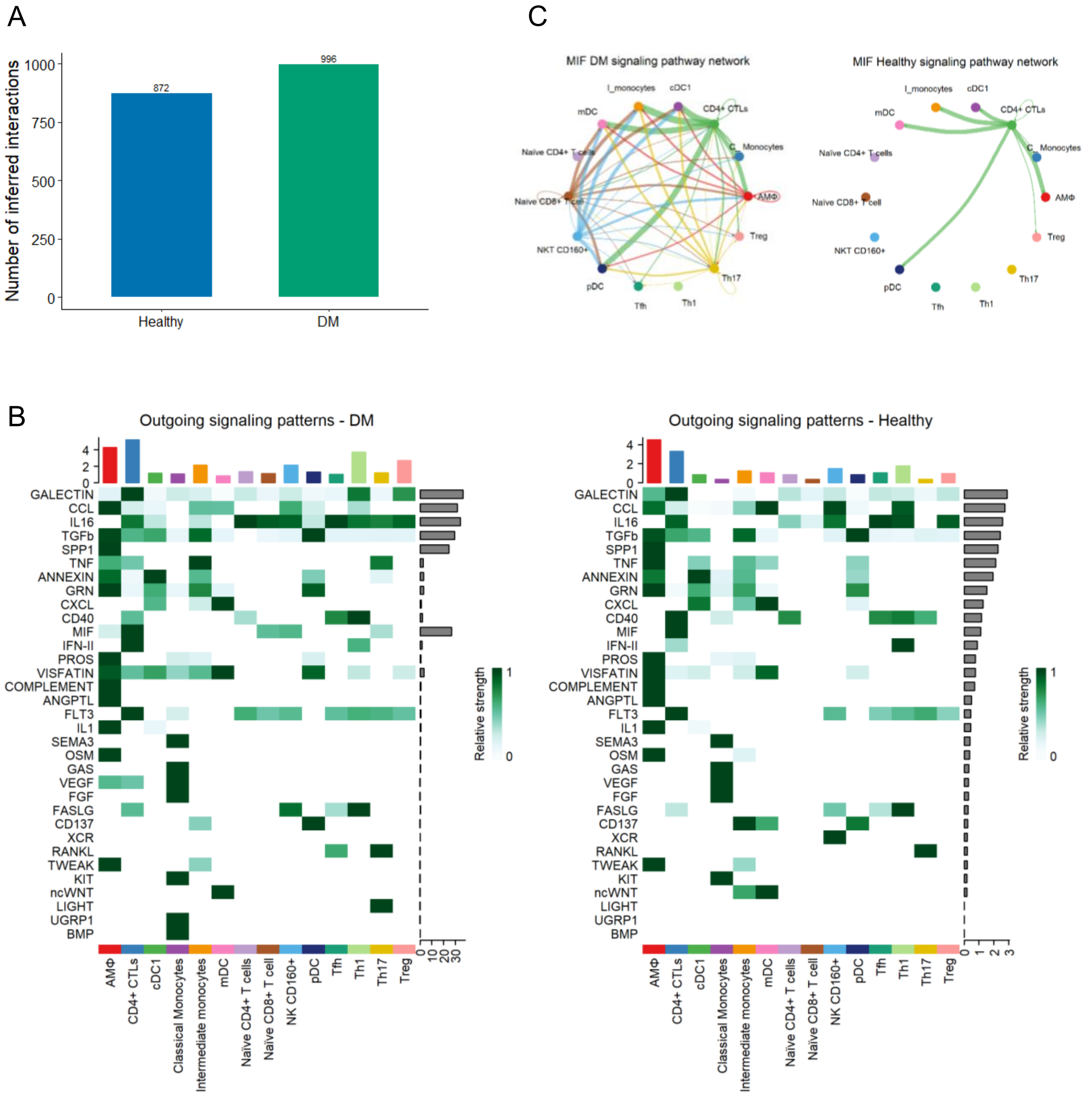


**Supplementary Figure S5:** (A) Bar plot showing the total number of inferred interactions for Healthy and DM groups. showing (B) Heatmap showing the contribution of outgoing signals to cell groups in DM and healthy groups. (C) MIF signalling network in DM and healthy groups. The thickness of the edge represents the strength of signaling.

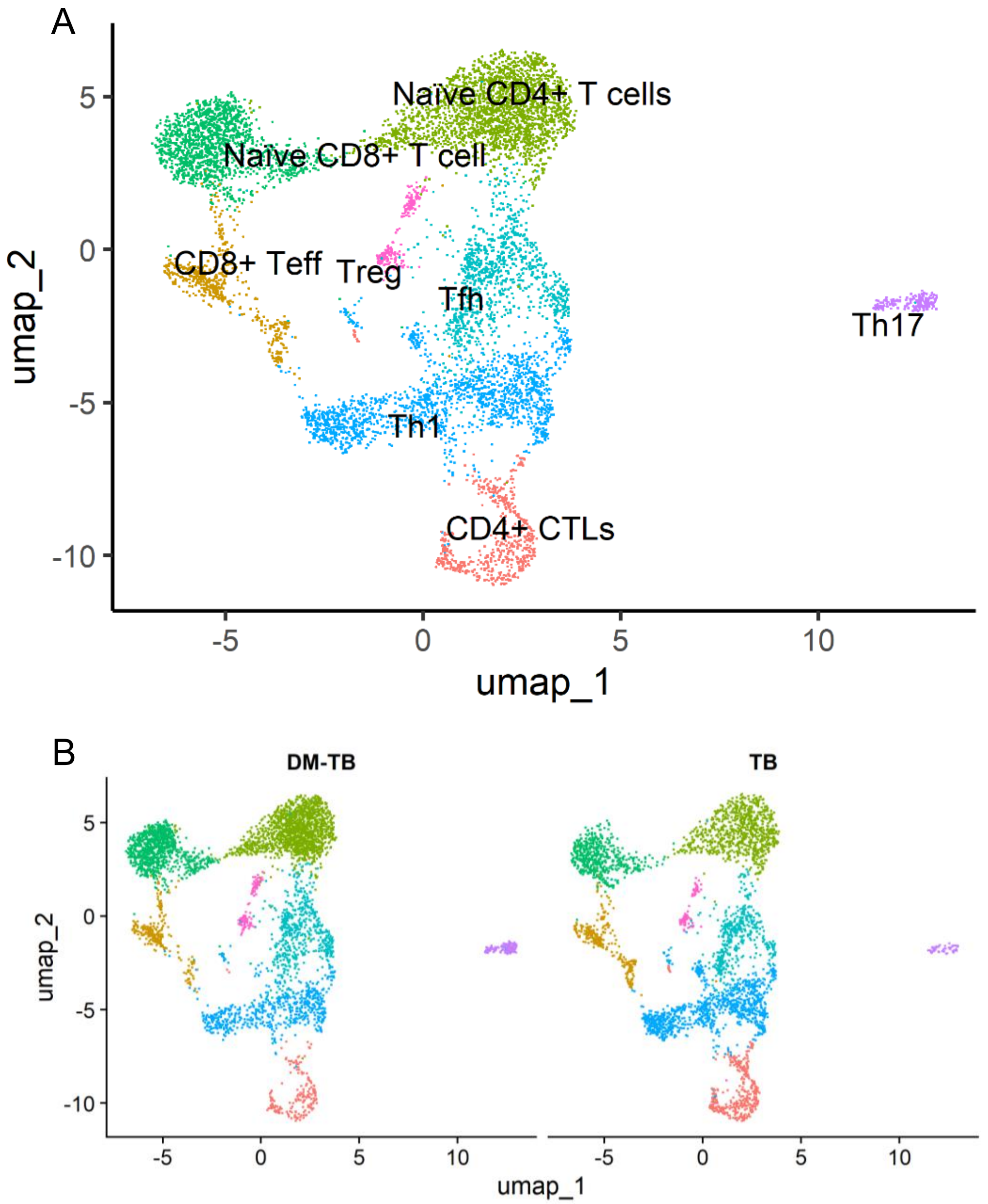


**Supplementary Figure S6:** Integrated (A) and split (B) T-cell UMAP for DM-TB and TB groups after sub setting for CD3+ cells. TB: *Mycobacterium tuberculosis* H37Rv infected control mice; DM-TB: Nicotinamide-Streptozotocin induced *Mycobacterium tuberculosis* H37Rv infected DM mice; CD8+ Teff: CD8+ effector T cells; Treg: Regulatory T cells; Tfh: Follicular helper T-cells; Th1: T helper cells type 1; CD4+ CTLs: CD4+ cytotoxic lymphocytes.

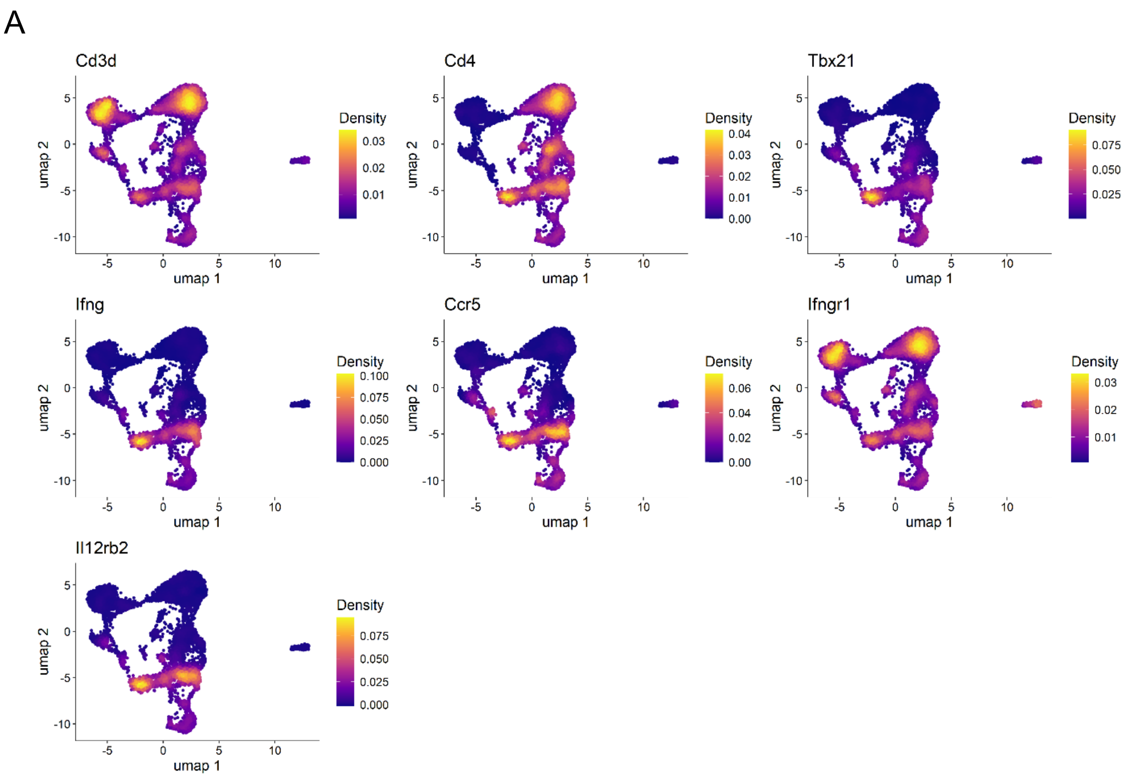


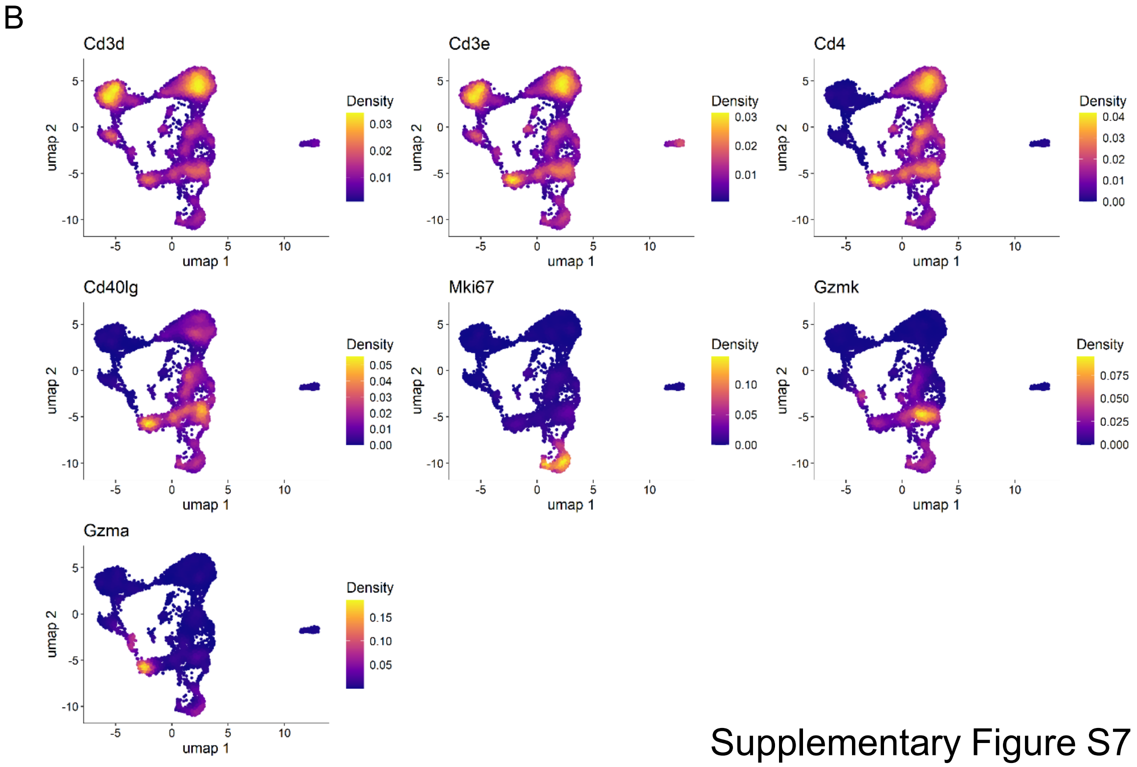


Supplementary Figure S7 continued

C

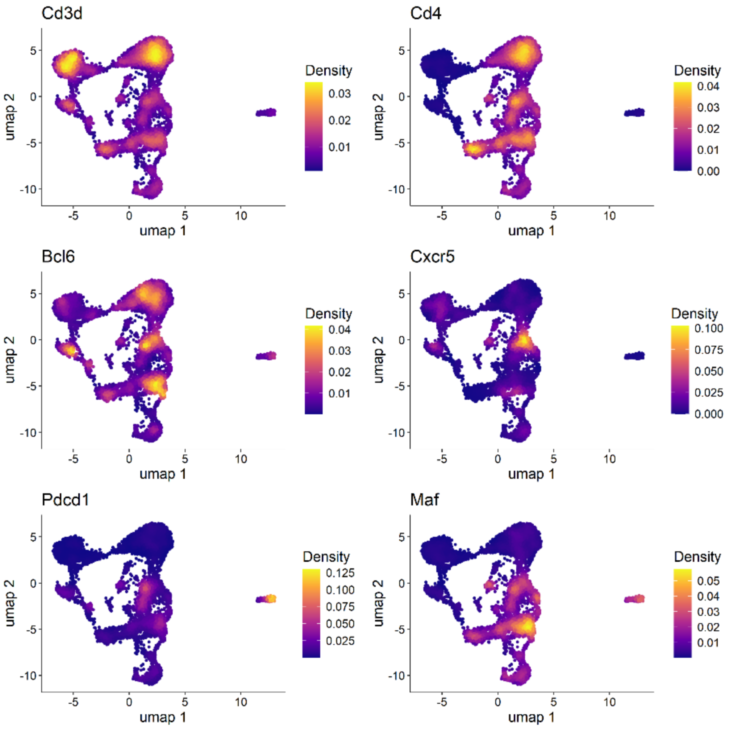


D

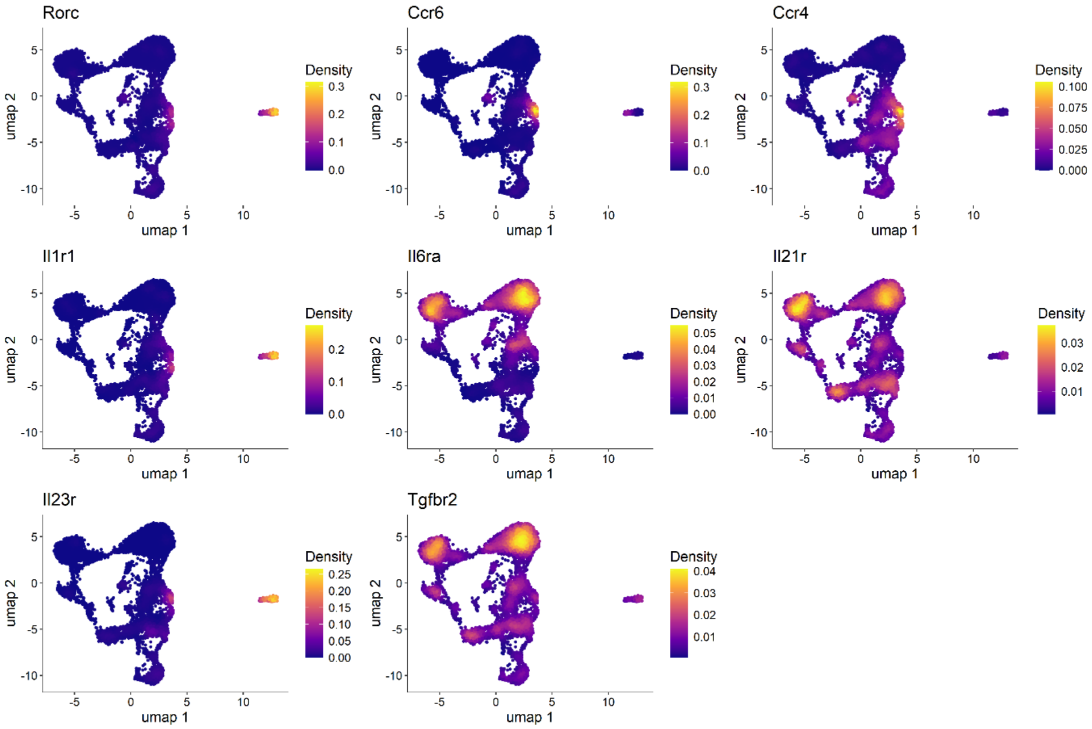


Supplementary Figure S7 continued

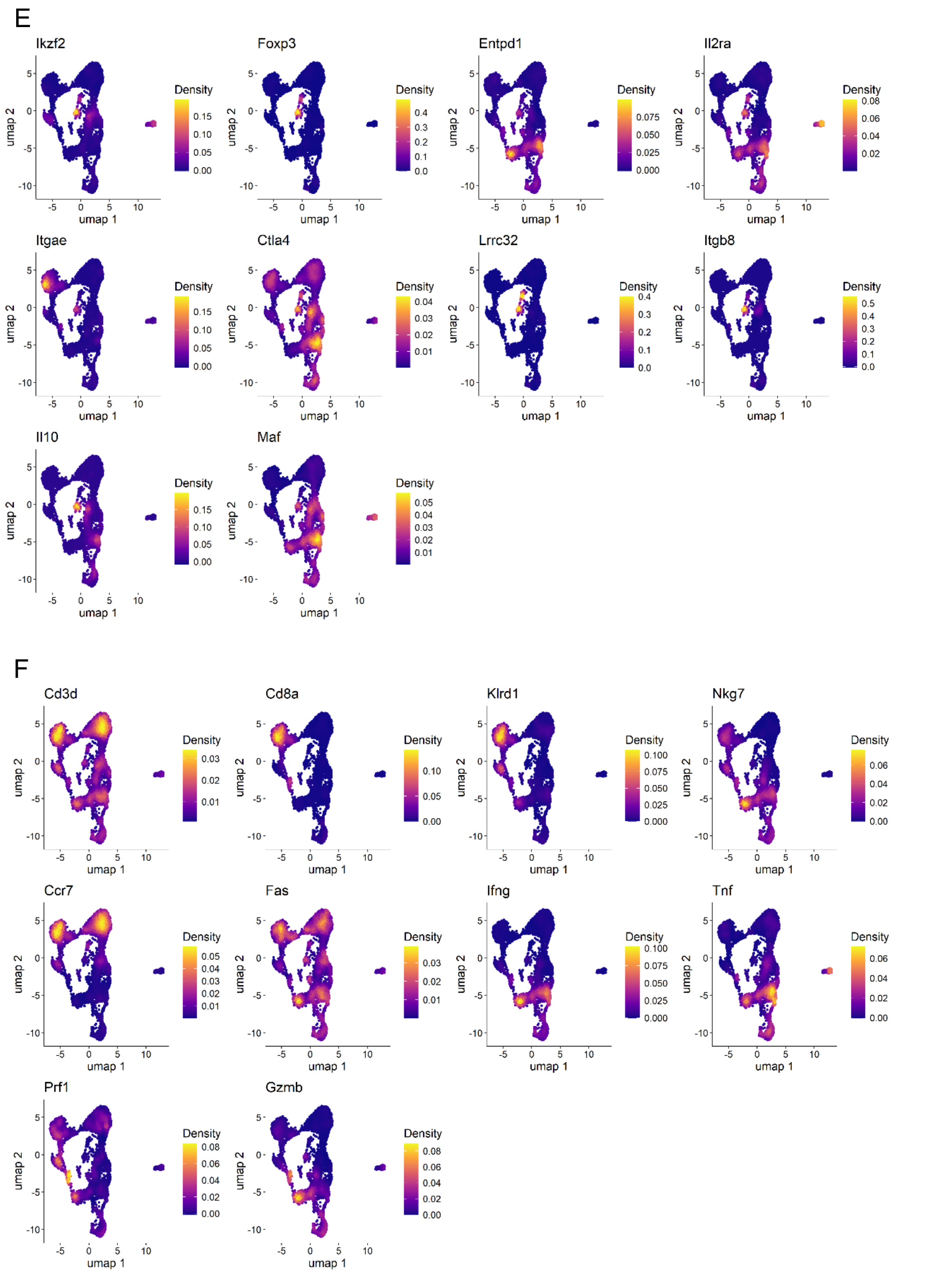


**Supplementary Figure S7:** Nebulosa plots showing expression density of canonical markers used for assigning (A) T-helper type 1, (B) CD4+ cytotoxic lymphocytes, (C) follicular helper T-cells, (D) T-helper type 17 cells, (E) regulatory T-cells and (F) CD8+ Effector T-cells.

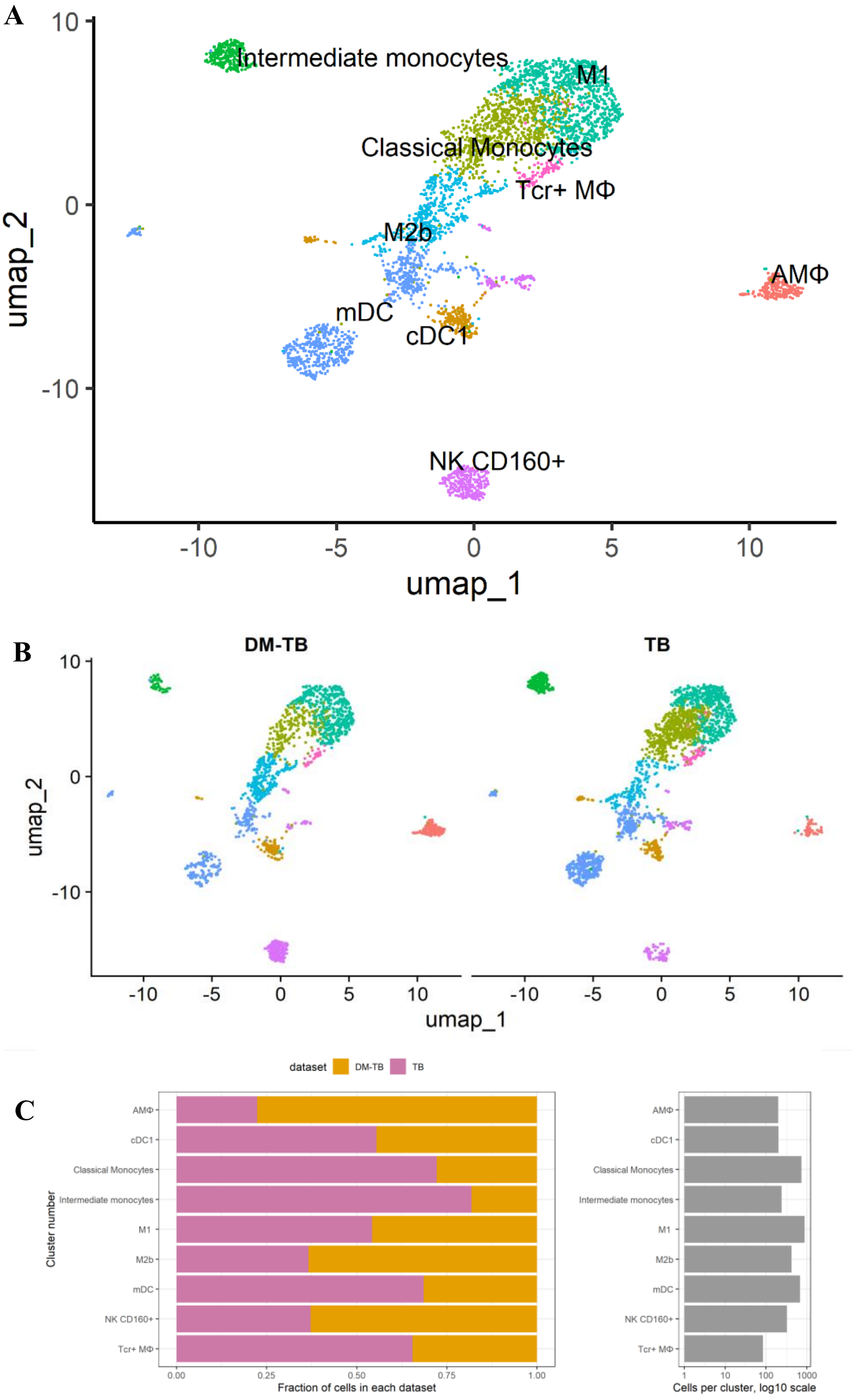


**Supplementary Figure S8:** Integrated (A), split (B) T-cell UMAP and (C) cluster metrics for DM-TB and TB groups after sub setting for Itgax+, CD14+, Itgam+ and CD3- cells. TB: Mycobacterium tuberculosis H37Rv infected control mice; DM-TB: Nicotinamide-Streptozotocin induced Mycobacterium tuberculosis H37Rv infected DM mice; M1: M1 macrophages; AMϕ: Alveolar macrophages; M2b: M2b macrophages; mDC: myeloid dendritic cells; cDC1: classical dendritic cells; Tcr+ Mϕ: T cell receptor+ macrophages; NK CD160+: Natural killer CD160+ cells.

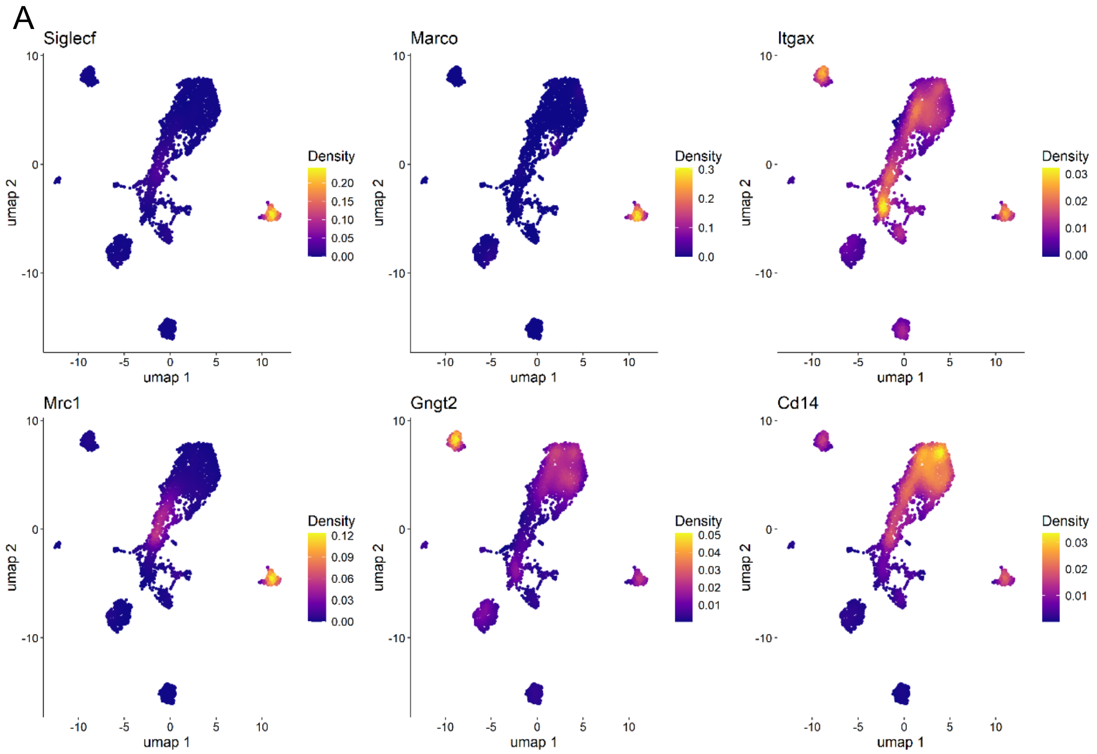


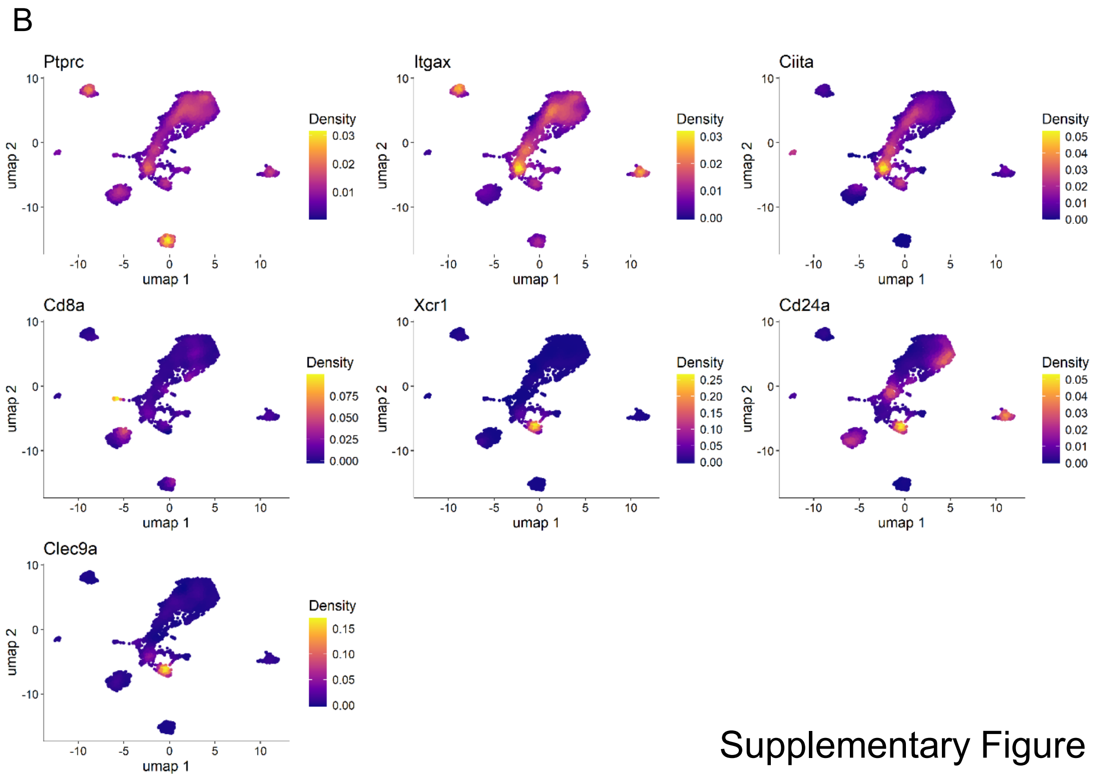


Supplementary Figure S9 continued

C

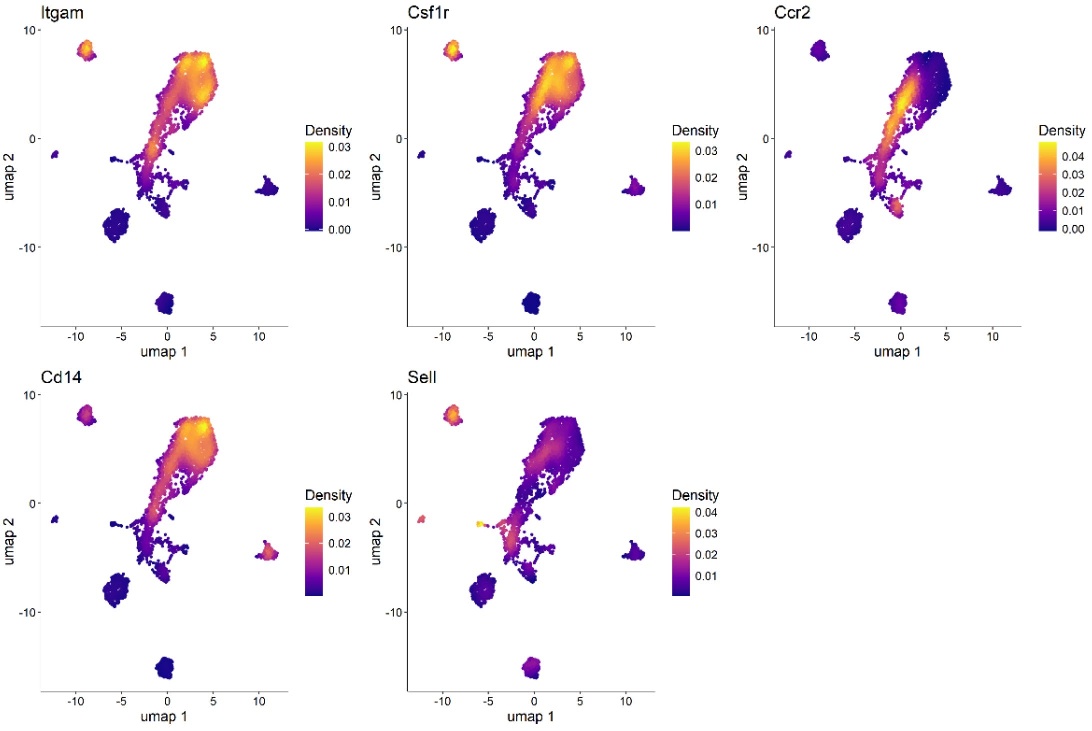


D

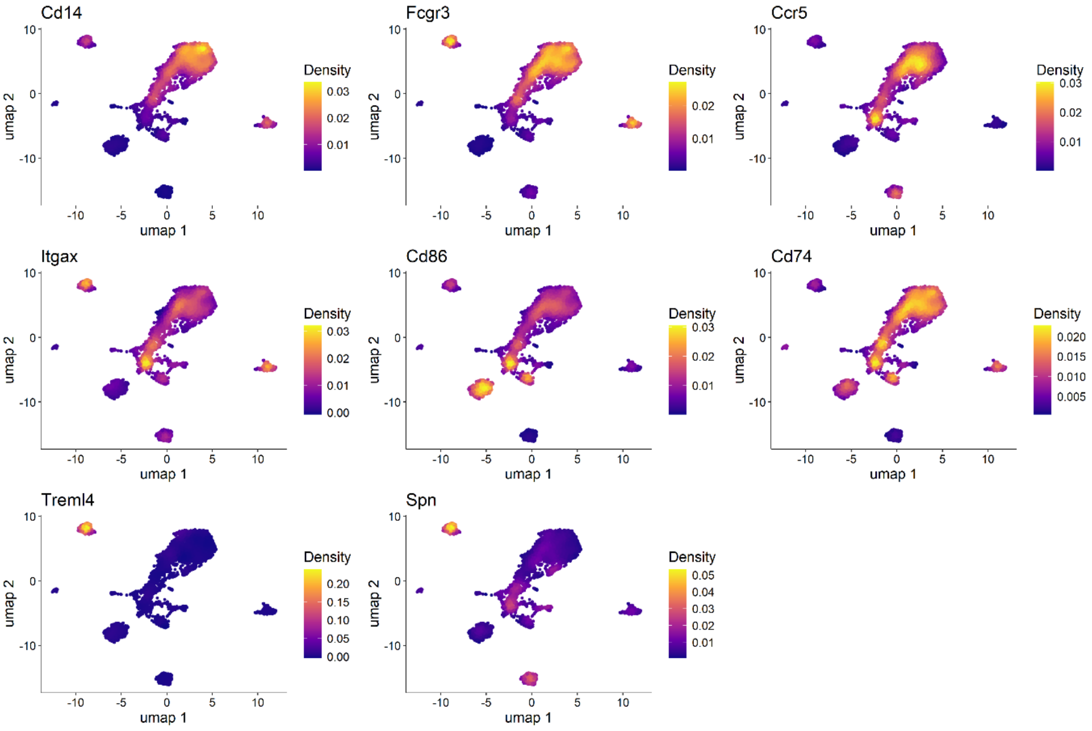


Supplementary Figure S9 continued

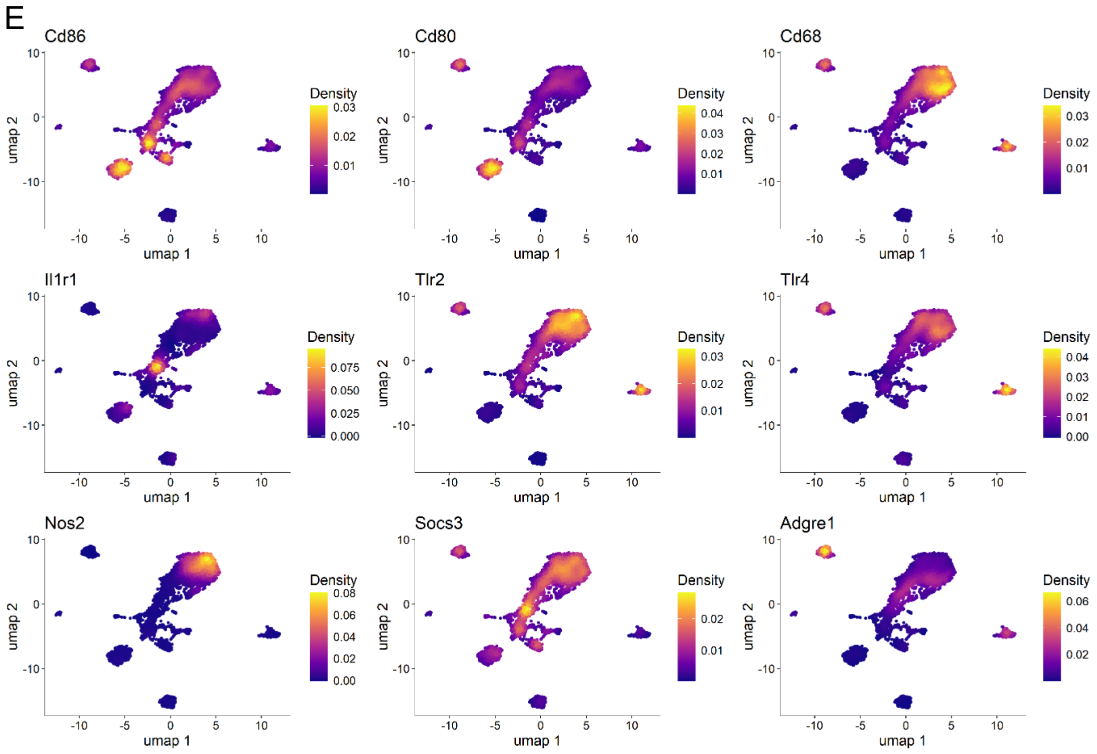


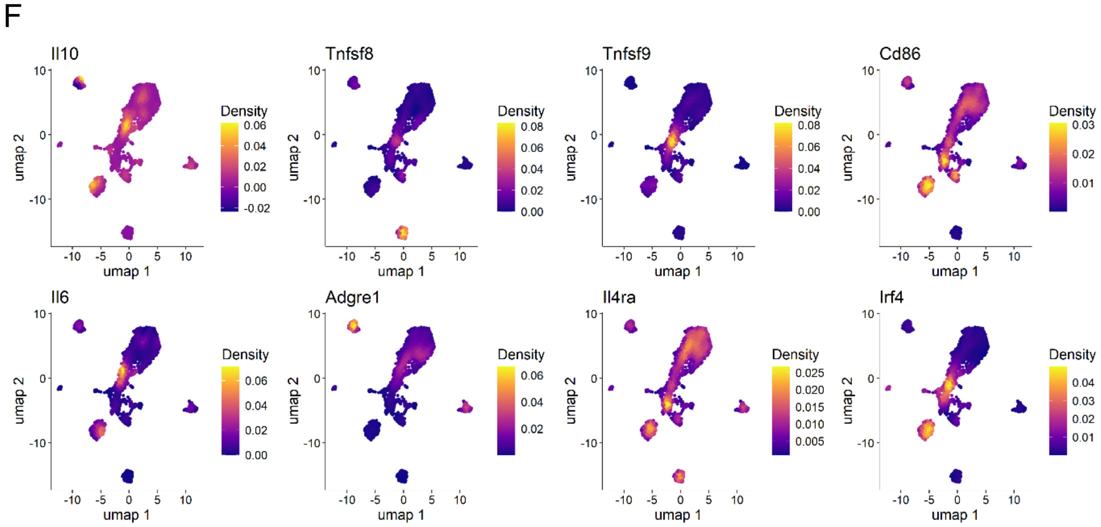


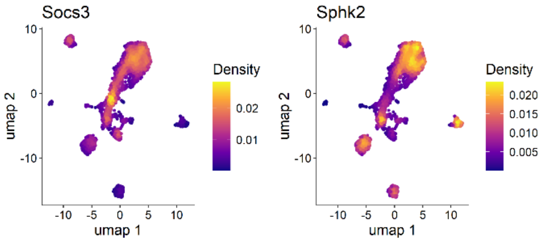


Supplementary Figure S9 continued

G

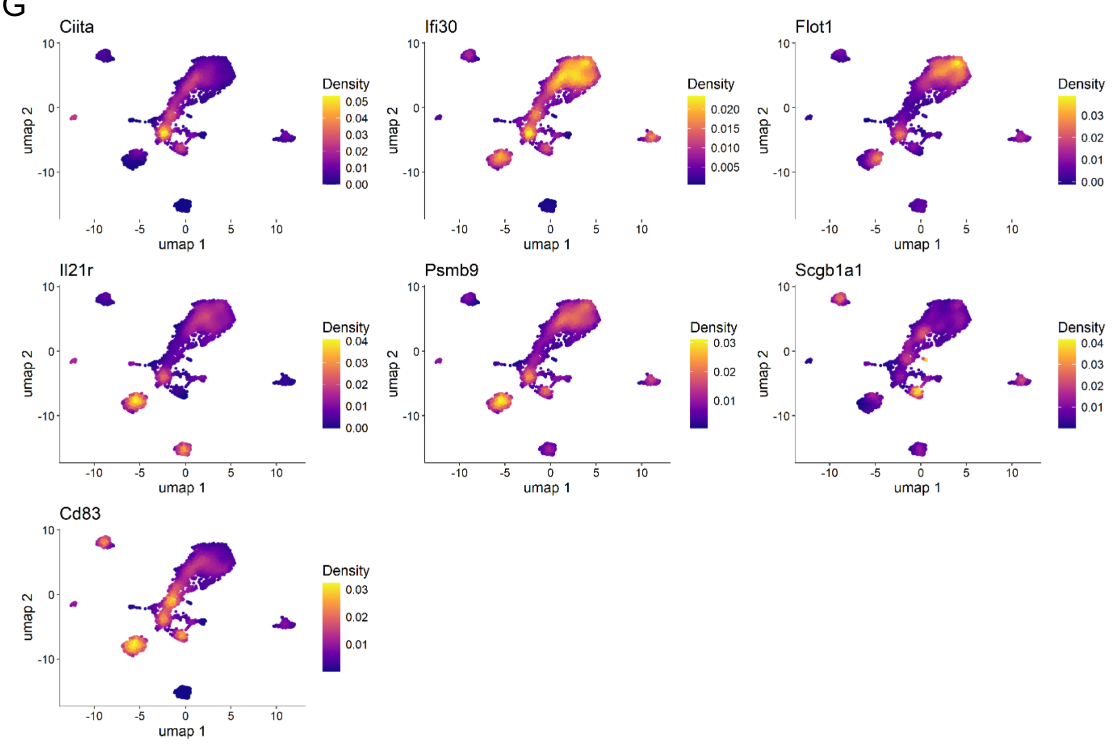


H

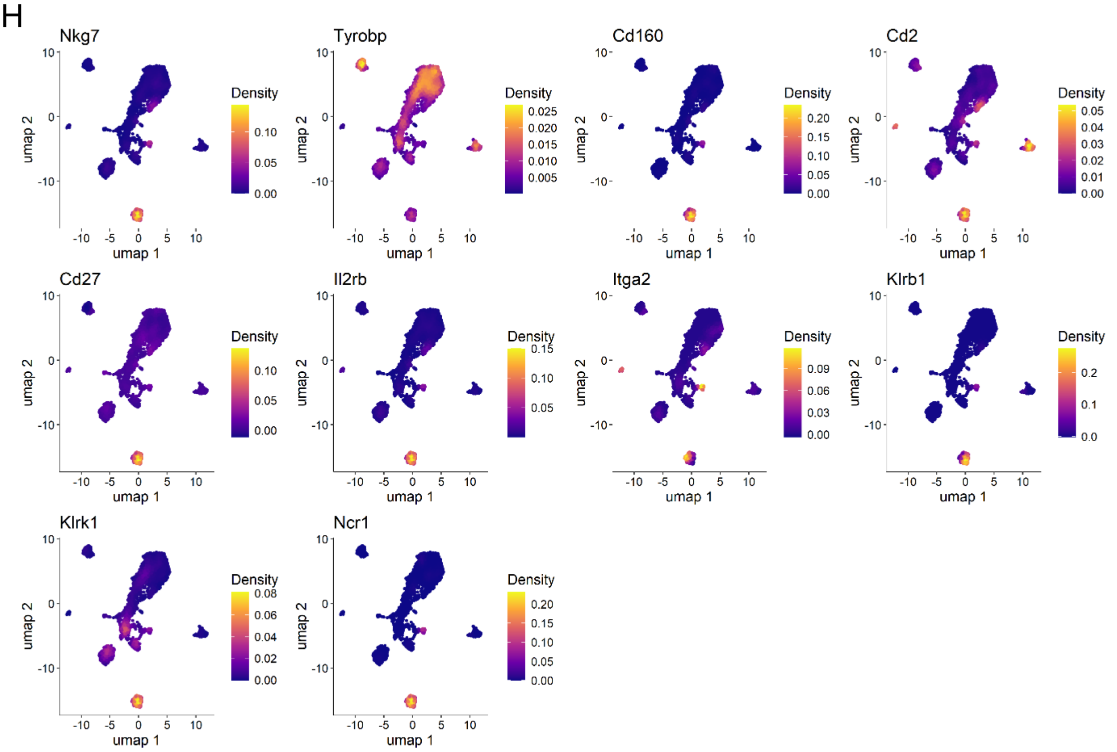


**Supplementary Figure S9:** Nebulosa plots showing expression density of canonical markers used for assigning (A) Alveolar macrophages, (B) classical dendritic cells, (C) classical monocytes, (D) intermediate monocytes, (E) M1 macrophages, (F) M2b macrophages, (G) myeloid dendritic cells and (H) Natural killer CD160+ cells.

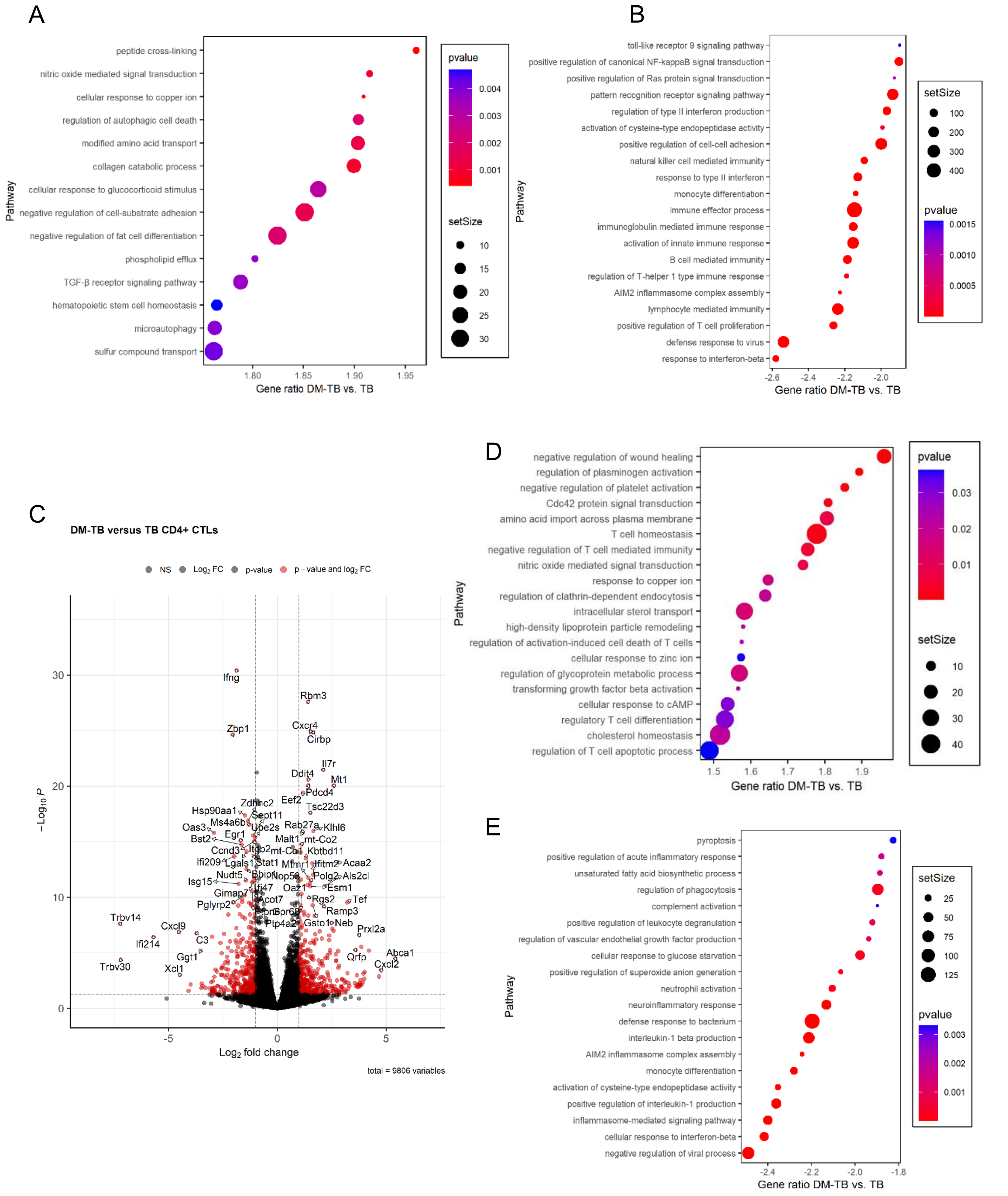


Supplementary Figure S10 continued

F

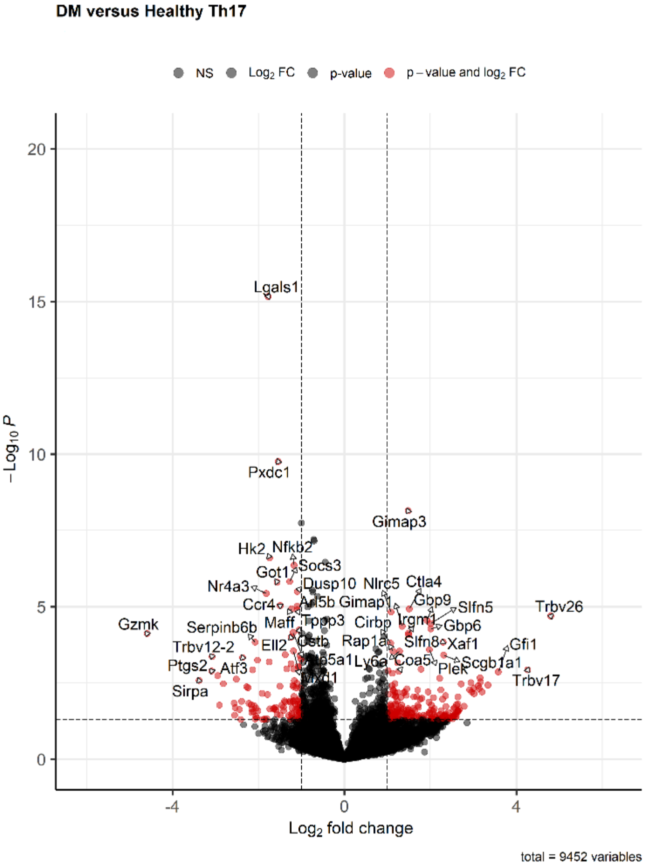


G

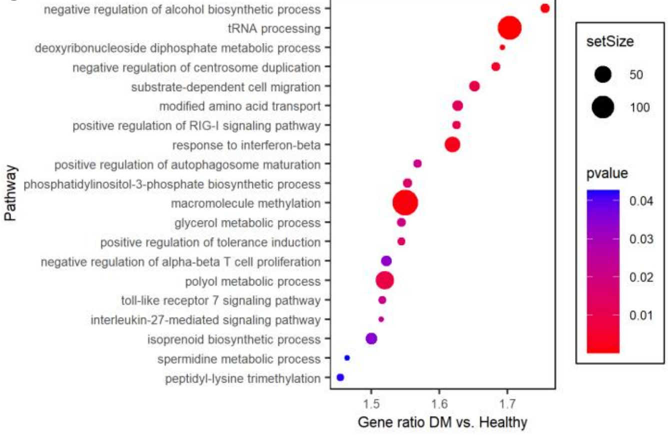


H

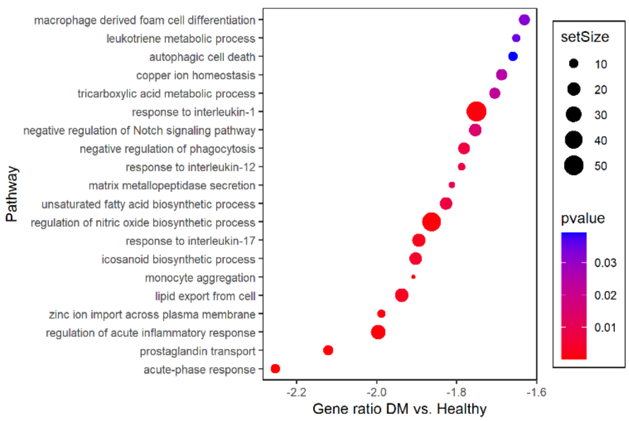


I

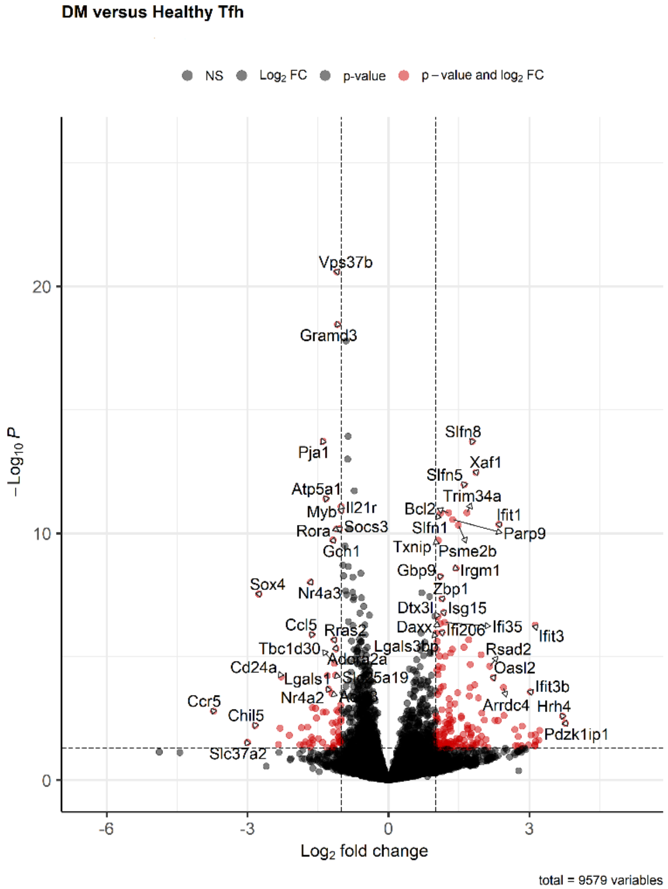


J

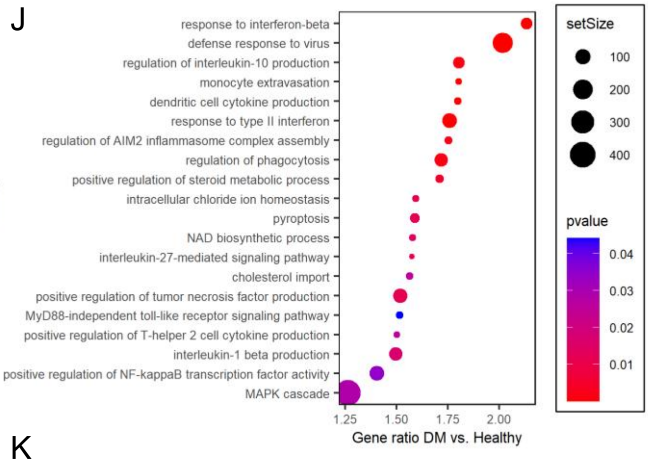


K

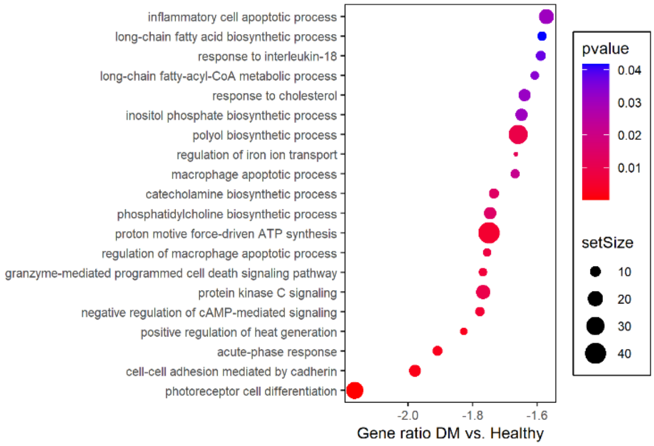


Supplementary Figure S10 continued

**Supplementary Figure S10:** GSEA for biological processes of M1 macrophages for (A) positively and (B) negatively enriched terms in DM-TB as compared to TB group. C) Volcano plot showing significantly upregulated/downregulated genes in CD4+ CTLs between DM-TB and TB mice. (D) Positively and (E) negatively enriched pathways in DM-TB CD4+ CTLs post GSEA analysis. (F) Volcano plot showing significantly upregulated/downregulated genes in Th17 cells between DM and healthy mice. Positively (G) and negatively (H) enriched pathways in DM naïve T-cells post GSEA analysis. (I) Volcano plot showing significantly upregulated/downregulated genes in Tfh cells between DM and healthy mice. Positively (J) and negatively (K) enriched pathways in DM naïve T-cells post GSEA analysis

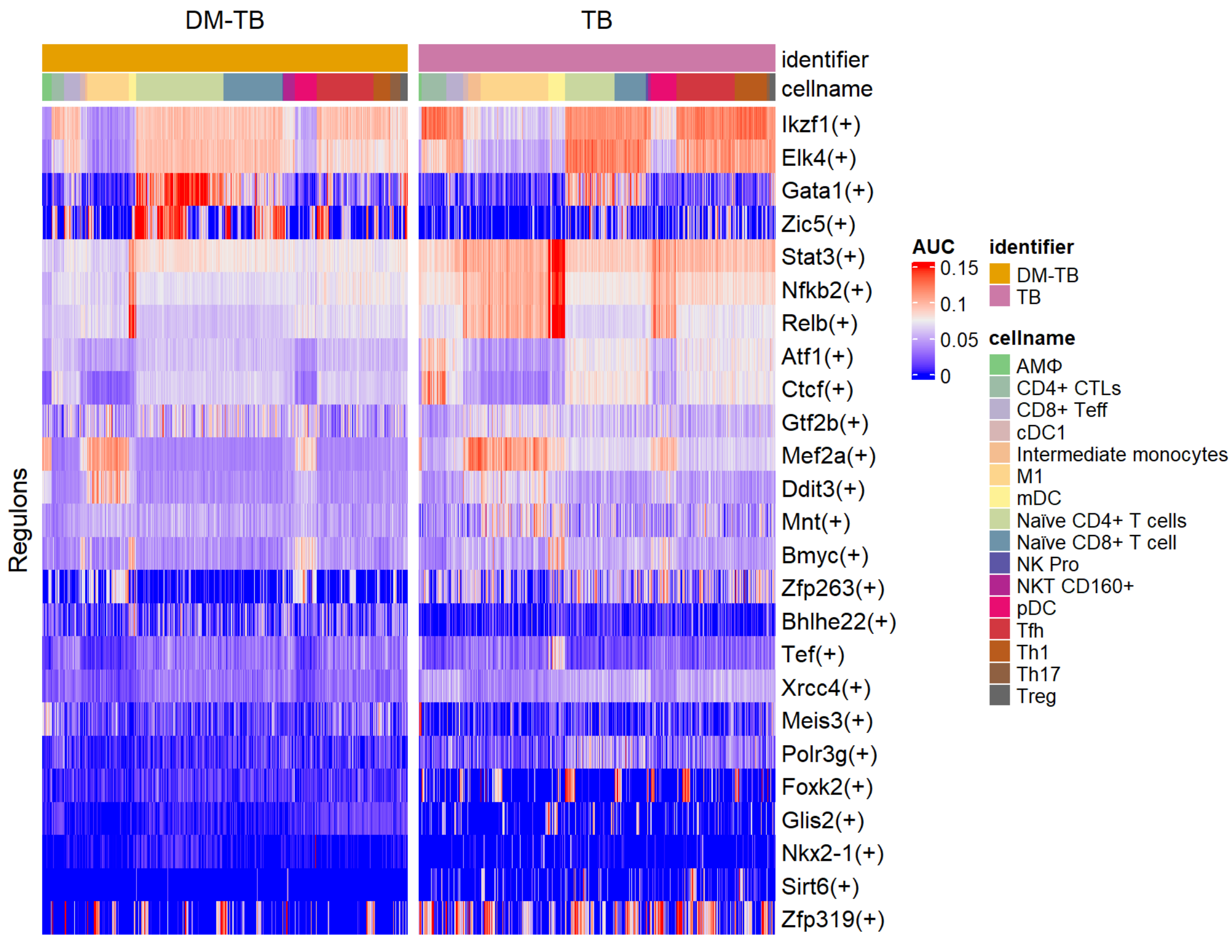


**Supplementary Figure S11:** Heat map showing differentially activated regulon for cell clusters in DM-TB and TB groups. Unpaired Wilcoxon rank-sum test was performed to compare the activity of each regulon between datasets (healthy and DM). Significant difference in regulon activity: p-value ≤ 0.05.

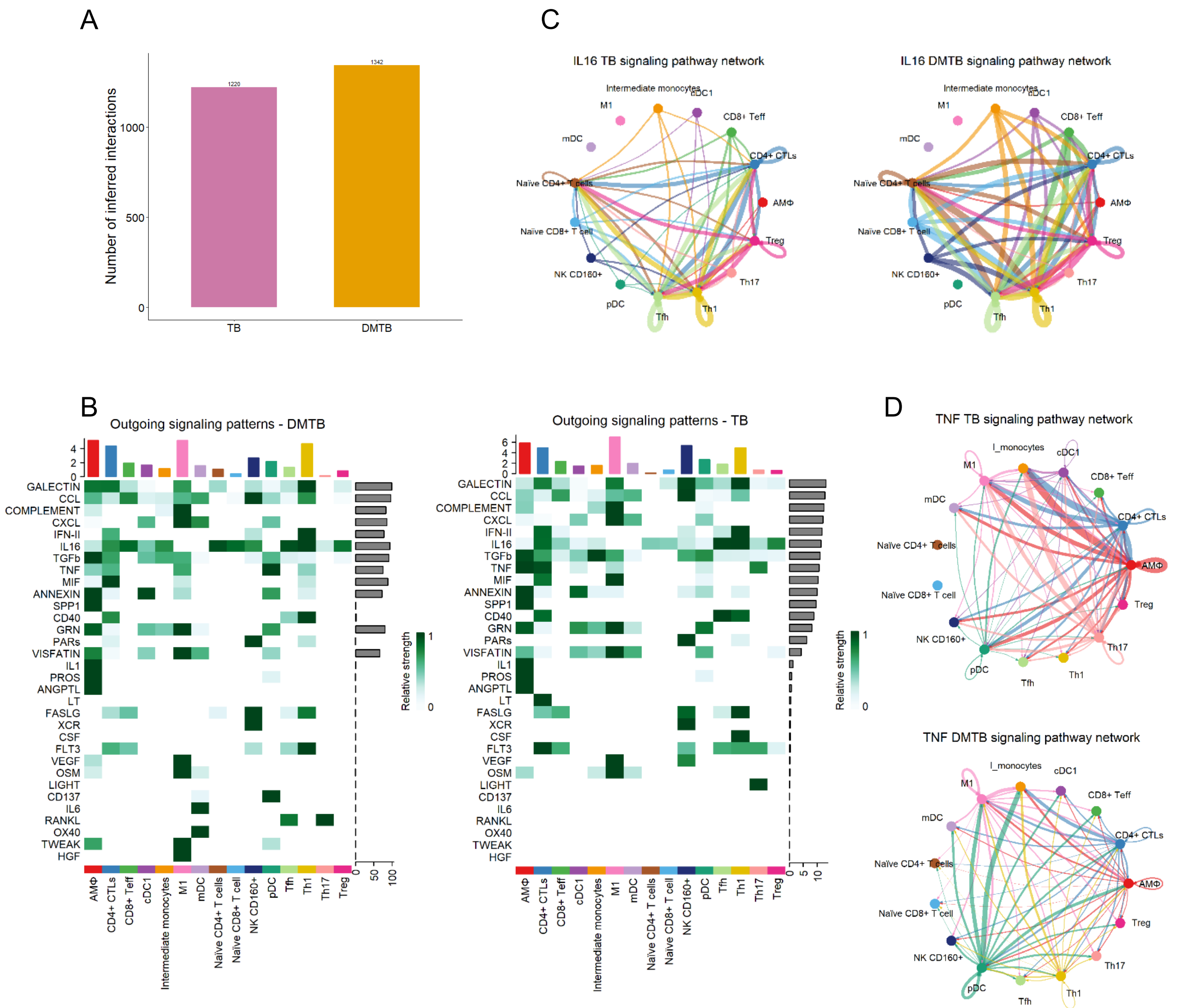


**Supplementary Figure S12:** Impaired anti-tuberculosis responses in diabetic mice following Mtb H37Rv infection. (A) Bar plot showing the total number of inferred interactions for TB and DM-TB groups. showing (B) Heatmap showing the contribution of outgoing signals to cell groups in DM-TB and TB groups. (C) IL-16 and (D) TNF signaling network in DM-TB and TB groups. The thickness of the edge represents the strength of signaling.
